# Targeted Next Generation Sequencing of environmental DNA improves detection and quantification of invasive European green crab (*Carcinus maenas*)

**DOI:** 10.1101/2021.03.01.433397

**Authors:** Kristen Marie Westfall, Thomas W. Therriault, Cathryn L. Abbott

## Abstract

In the northeast Pacific Ocean there is high interest in developing eDNA-based survey methods to aid management of invasive populations of European green crab (*Carcinus maenas*). Expected benefits are improved sensitivity for early detection of secondary spread and quantification of abundances to assess the outcome of eradication efforts. A new eDNA-based approach we term ‘Targeted Next Generation Sequencing (tNGS)’ is introduced here and shown to improve detection relative to qPCR at low eDNA concentrations, as is characteristic of founding or spreading populations. tNGS is based on the premise that the number of NGS reads from non-normalized (i.e. equal volumes) targeted PCR amplicons will approximate the starting DNA amount. Standard DNA concentrations that were 10-to 100-times lower than the qPCR limit of detection returned significant numbers of sequencing reads, which in our field assessments translated to a 7% - 10% increase in crab detection probability from tNGS relative to qPCR at low abundances. We also found that eDNA concentration was highly correlated with crab abundance, as measured from traditional trapping methods, for both assays; however, tNGS data had greater precision and less error than qPCR. When partitioning the sources of variation in each assay we identified greater between-site variability for tNGS relative to qPCR, suggesting the former may offer more power for detecting spatial variation in eDNA concentration. When applying this assay in management programs, we suggest including a panel of eDNA samples from sites with trapping data as standards to estimate relative abundance at sites with no a priori information. Results presented here indicate the tNGS approach has great promise for surveillance of green crab and could easily be adopted for surveillance of any species of high interest to management, including endangered species, new incursions of invasive species, and species with low eDNA shedding rates. Pros and cons of this approach compared to qPCR are discussed.

## 1.0 Introduction

Population abundance and distribution are foundational tenets of the ecological study (Krebs, 1972) and management of species (Clement et al., 2015; Haines et al., 2013; Jones, 2011; Yin & He, 2014), but can be difficult to measure for endangered species, secretive or rare species, or in remote or isolated environments. Accurate, rapid, and low-cost quantitative methods for assessing these population metrics would benefit many branches of resource and conservation management by providing essential data for regulatory decisions; for example, as related to population status updates and recovery plans (or benchmarks) (Haines et al., 2013). Invasive species often are considered rare in the very early stages of the invasion process. Detection of invasive species during this phase is paramount for quick implementation of control efforts (Harvey et al., 2009; Howald et al., 2007; Lodge et al., 2006) and for judging the efficacy of such actions (Davis et al., 2016). Validating lower-cost and less intensive methods for these population assessments, especially for rare species, is an essential step forward for effective management of scarce species.

Environmental (e)DNA is extra-organismal DNA obtained directly from the environment, deriving from shed cells, faeces, mucous, gametes, carcasses, and myriad other sources (Sassoubre et al., 2016; Thomsen & Willerslev, 2015). ‘Community DNA’ is the term given to a mixture of organisms isolated from an environmental sample (Deiner et al., 2017). Species detection from eDNA has particular advantages; environmental samples are relatively easy to collect, species can be detected low densities, and species can be detected throughout their life cycle (Deiner et al., 2017; Furlan et al., 2019; Taberlet et al., 2012). Rare, endangered (Franklin et al., 2019; Hunter et al., 2018; Laramie et al., 2015; Thomsen et al., 2012), and invasive (Holman et al., 2019; Westfall et al., 2019) species have been identified from eDNA across a range of habitats (terrestrial, marine, freshwater).

The application of eDNA is generally limited to presence/absence surveys, but eDNA has shown to be positively correlated with species abundance in laboratory and field-based surveys (Doi et al., 2017; Knudsen et al., 2019; Lacoursière-Roussel et al., 2016; Lehman et al., 2019; Shelton et al., 2019). High variation in quantitative results from field surveys and a lack of rigorous validation in controlled laboratory settings have lead to considerable scrutiny of these methods (Mauvisseau, Burian, et al., 2019). However, eDNA is intrinsically variable both spatially and temporally within the same environment due to water movements (linear flow in rivers or tidal flow in the ocean), seasonal reproductive stages, or different DNA shedding rates across species (Forsström & Vasemägi, 2016).

Field census surveys using traditional methods are subject to the same rules of nature and have additional biases that affect ‘catchability’ across species (Duncombe & Therriault, 2017). The value of eDNA lies in it’s ease of sample collection, lower cost, and non-invasive sampling. It has also repeatedly detected species in some of the most important cases where traditional methods failed, eg. highly endangered and scarce species (Franklin et al., 2019; Laramie et al., 2015; Mauvisseau, Davy-Bowker, et al., 2019) or at the first onset of a potentially damaging bioinvasion (Dougherty et al., 2016; Holman et al., 2019).

Quantitative eDNA applications have been successfully compared with traditional survey methods for species at risk (Lehman et al., 2019; Shelton et al., 2019), invasive species (Thomas et al., 2019), and fisheries stock assessments (Doi et al., 2017; Knudsen et al., 2019; Lacoursière-Roussel et al., 2018). In general, eDNA availability of a focal species is positively correlated with the abundance of that species in the vicinity (Doi et al., 2017; Knudsen et al., 2019; Lacoursière-Roussel et al., 2016; Lehman et al., 2019; Shelton et al., 2019). The decay rate of eDNA has been experimentally observed at hours to days and the geographic area which it may represent is highly variable among environments (i.e. fresh vs. saltwater, lentic vs. lotic) and sample types (i.e. sediment vs. water). The correlation between eDNA concentration and abundance is therefore highly variable among species and environments, and is vulnerable to seasonal variation (Doi et al., 2017), breeding characteristics (Buxton et al., 2017), and hydrographic variation (Carraro et al., 2018; Tillotson et al., 2018). Furthermore, the precision of abundance estimates is known to correspond with eDNA concentration (i.e. low precision at low concentration) and thus is sensitive to sampling characteristics like the number of replicate subsamples taken from a site or at each sampling period (Doi et al., 2017; Pilliod et al., 2013).

Decreased precision at low eDNA concentrations is also linked with intrinsic technical limitations of quantitative PCR methods such as quantitative (q)PCR or digital droplet (d)PCR, the former being the most common method for targeted species detection and quantification. False positive fluorescent background signals (Strain et al., 2013) and contamination from non-specific amplification can lead to over-estimation of eDNA at low concentrations (Hunter et al., 2018; Langlois et al., 2020), such as would be expected from rare species eDNA. Stochastic variation of molecule distribution in an aqueous sample and the propensity of DNA to adsorb onto plastic tube walls and pipette tips are known to affect the proportion of positive results from low template concentrations (Ellison et al., 2006). The Minimum Information for Publication of Quantitative Real-Time PCR Experiments (MIQE) guidelines define the limit of detection (LOD) as the ‘lowest concentration at which 95% of positive samples are detected’ (Bustin et al., 2019), but some eDNA qPCR assays set the LOD at the concentration for which 50% or fewer replicates are positive (Biggs et al., 2015; Harper et al., 2018; Roux et al., 2020). In an effort to standardize reporting guidelines eDNA qPCR assays, Klymus et al. (2020) (Klymus et al., 2020) suggested that LOD be defined as for MIQE for quantitative applications, but also suggested that qualitative detections can be called at less than 40 cycles.

Here we propose a new quantitative eDNA assay that circumvents the shortcomings of fluorescence-based quantitative technologies in the application to low copy number eDNA surveys. Termed ‘targeted NGS’ (tNGS), PCR amplicons from targeted primers (i.e. those designed for qPCR assays) are directly sequenced on the Illumina platform along with standardized DNA much like a traditional qPCR assay. The premise is that when eDNA is amplified in a single step PCR protocol and products sequenced at equal volumes without normalization, the number of sequencing reads will have a strong positive correlation with eDNA concentration and, therefore, with species abundance. The primers are developed under the same MIQE guidelines for qPCR, including meeting the standards for specificity and sensitivity. Direct sequencing of PCR amplicons has many potential benefits over traditional quantitative PCR assays; there is no background fluorescence to infer false positives at low template concentrations, sequences can identify non-target species, and potentially quantify abundance.

Regardless of whether the goal of an eDNA quantitative assay is to complement or replace an existing traditional method for detection and estimating abundance, it is important to empirically demonstrate that eDNA can provide a similar level of information. If eDNA can be used as a first line of detection and quantification, then resources for more labour intensive surveys can be targeted to specific areas. The current study applies tNGS in an ideal test case of the invasive European green crab (*Carcinus maenas*; Linnaeus, 1758) on the Pacific coast of Canada. Invasive green crab have established throughout Barkley Sound (Vancouver Island) with extreme patchiness in population densities that have been catalogued by census surveys since 2004. Previous research developed a qPCR assay to complement existing census surveys and implement an early detection system for secondary spread into neighbouring regions (Roux et al., 2020). In the current study we aim to directly compare results from the qPCR assay of Roux et al. (2020) with tNGS to answer the following key questions: (1) do tNGS and qPCR have similar probabilities of detecting green crab?; (2) do spatial patterns of eDNA concentration correlate with crab abundance based on trapping, and are the assays equally powerful at detecting spatial variation?; and (3) what field sampling considerations maximize detection? We conclude with suggestions on how to best apply this method in management programs.

## 2.0 Materials & Methods

### 2.1 Study site and green crab trapping

Green crab trapping, water and plankton sampling were conducted in Barkley Sound in August 2017, Vancouver Island, Canada (Fig. 1 and Supplementary Table S1) at sites with regular trapping data since 2004. Based on these existing trapping data (not shown), three higher abundance sites (Pipestem Inlet, Mayne Bay, and Hillier Island) and two lower abundance sites (San Mateo Bay, and Ritherdon Bay) were selected (Fig. 1). Consistent with established green crab survey methods (i.e., Gillespie et al., 2007), three sets of six traps were baited and deployed in the intertidal/shallow subtidal zone at each site. Traps were retrieved after ∼24 hours and green crab were sexed and enumerated. Catch per unit effort (CPUE) was calculated as the number of green crab caught divided by the number of traps counted. The number of traps counted varied across sites due to loss from black bear activity.

**Fig. 1.**
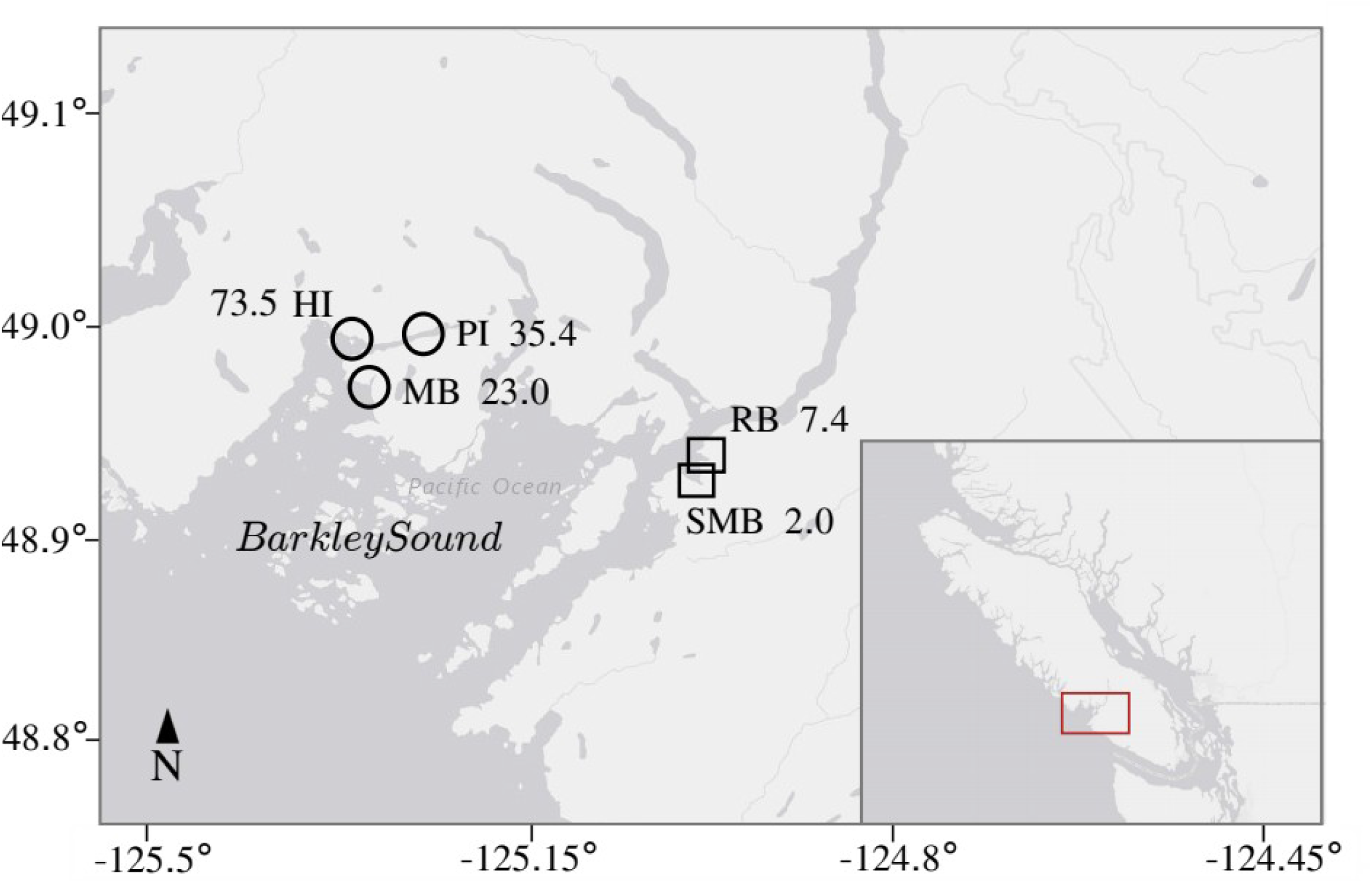
Map of green crab sample sites in Barkley Sound with green crab catch per unit effort indicated for each site. Large map is red inset of smaller map, which shows Vancouver Island, BC, Canada. Sites marked by an open circle are high density sites Pipestem Inlet (PI) and Mayne Bay (MB). Sites marked by an open square are low density sites Ritherden Bay (RB) and San Mateo Bay (SMB). Latitude on x-axis and longitude on y-axis, both in degree decimal format.

### 2.2 DNA sampling and extraction

Environmental DNA from filtered sea water and community DNA from zooplankton samples were compared to assess relative efficacy for green crab detection and quantification. Water sampling proceeded by taking six site replicates of 1L surface water on each of two days, totalling 12 subsamples per site (six per day). To minimize inflation of the eDNA signal due to trapping activities, water sampling on the first day was done prior to any green crab trapping gear being deployed and on day two was done prior to the gear being retrieved. Taking two sets of samples ∼24 hours apart was implemented to assess variation in resulting eDNA concentrations. All sample containers were sterilized prior to use by submerging into a 1% sodium hypochlorite solution for one minute and rinsing thoroughly with tap water. One vertical zooplankton tow of 10m depth was conducted at each site during trap deployment using a Bongo style net with 350 µM mesh size. Zooplankton samples were immediately fixed in 70% ethanol and stored at 4°C until DNA extraction. Plankton nets and accompanying equipment were sterilized between sites by submerging into a 1% sodium hypochlorite solution for one minute and rinsing thoroughly with sea water. Field blanks of 1L tap water were employed to test for eDNA contamination at this source.

On the same day as collection, water samples and field blanks were filtered through 47mm diameter cellulose-nitrate membrane filters with a 1µm pore size (Whatman) using a vacuum pump. All filtration equipment was sterilized using a 1% sodium hypochlorite solution for at least one minute and rinsed in tap water prior to filtering each sample. Equipment blank samples of 1L tap water were filtered after each equipment sterilization as a control. Filters were stored in 1.5 mL tubes with tissue lysis buffer ATL (QIAGEN) and proteinase K (QIAGEN). All filters were incubated the day of sampling for 10 hours at 37°C and stored at 4°C until DNA extraction.

Extraction of eDNA was performed directly from the digest in the 1.5mL tubes with filters using the standard protocol for animal tissue of the DNeasy Blood & Tissue kit (QIAGEN). Extraction of community DNA from zooplankton samples proceeded by volumetrically splitting the sample into two equal parts, decanting the ethanol, rinsing and drying the samples for a short period, homogenizing the tissue with a mortar and pestle and taking 2mL of homogenate. All equipment was sterilized between samples using a 1% sodium hypochlorite solution for at least one minute and rinsed in tap water. Blank controls of tap water were extracted in each eDNA extraction session.

### 2.3 Laboratory protocols for green crab assays

#### 2.3.1 Data sets

Two approaches were used to process water sample replicates, thus generating two eDNA data sets that were used to compare how field sample replication impacted green crab detection and quantification. The first data set consisted of all eDNA water sample replicates treated separately (herein *replicate*) for a total of 48 eDNA samples (6 site replicates x 4 sites x 2 days). Note that 4 sites were analyzed for the *replicate* data set. A second eDNA data set was created by pooling the six eDNA water sample replicates from each site on each day into one sample, which resulted in each site thus having one pooled eDNA sample per day (herein *site*) for a total of 10 eDNA samples (5 sites x 2 days). The zooplankton DNA data set consisted of a total of five samples (1 day x 5 sites), each of which was treated separately.

#### 2.3.2 Quantitative PCR

Quantitative PCR was performed for *replicate* samples by Roux et al. (2020) and was performed here for *site* and zooplankton samples following the same protocol exactly and in the same lab. Three qPCR technical replicates were performed for all samples. In cases where only one qPCR technical replicate returned a C_t_ value, that value was used to represent the whole sample; in cases where more than one qPCR replicate returned a C_t_ value, these were averaged to represent the whole sample. These whole sample level C_t_ values were then used to calculate eDNA concentration for that sample. A sample was labelled non-detect if all technical replicates did not to return a C_t_ value. Two alternative methods to deal with non-detects were compared in the modelling analyses described below: (1) remove them from the data set; or (2) assign them a C_t_ value of 40 (i.e. the number of PCR cycles performed). Since there were no differences in model output conclusions from both methods, the most parsimonious method of full removal was used for final analyses.

#### 2.3.3 Targeted NGS

Sequencing was performed by amplifying in a single-step PCR using the same species-specific primers as used for qPCR, but within longer fusion primers that also included the Illumina sequencing tags (Supplementary Table S2). Single-step PCR eliminates product mass variation due to ligation efficiencies, bead recovery, and other steps involved in Illumina library preparation and normalization. Fusion primer design and pairing scheme roughly followed that from Elbrecht & Leese (2015) with an additional pentamer barcode to aid in demultiplexing (Supplementary Table S2). High positional diversity was ensured by: (1) 15% PhiX spike; (2) shifting the base composition of any given position by adding zero to four bases to the primers; and (3) simultaneously sequencing forward and reverse primers (Elbrecht & Leese, 2015) (Supplementary Table S2). PCR reactions were performed in triplicate with 3 µL template in a 25 µL reaction with final concentrations: 1µM each primer, 2mM dNTP, 1 mg/mL BSA, 2.5mM MgCl2, 1X buffer, and 2U Amplitaq Gold. Cycling conditions were 95°C for 10 min followed by 40 cycles of 95°C for 10 sec, 60°C for 1 min. PCR replicates of all water and zooplankton samples, blanks, and standard curve concentrations (see next paragraph) were pooled at equal volumes into two sequencing pools and cleaned twice with SPRI beads at 0.9x ratio. Libraries were quantified using a SYBR-Green fluorometric assay read on the Tecan Infinite Pro (Alam et al., 2017) and 15 pmoles of each library were loaded on two runs of MiSeq v2 300 cycles (150bp paired end).

Raw fastq files were trimmed of adapter sequence, low quality regions (phred33 score < 2), and N’s using Adapter Removal v2 (Schubert et al., 2016). OBITools v 1.2.11 (Boyer et al., 2016) was used to merge paired-end reads (‘illuminapairedend, minimum alignment score 40). JAMP v0.67 (github.com/VascoElbrecht/JAMP) was used to demultiplex the samples. All samples were screened for green crab reads using BLAST+ with a custom database of all green crab COI sequences available in the NCBI non-redundant nucleotide database (downloaded Feb 27, 2019) (Camacho et al., 2009). All non-target sequences were screened by matching to the PhiX genome used by Illumina (Genbank Accession NC001422) with BLAST+.

Standard curves were included in each run to quantify the relationship between starting DNA concentration and the number of raw reads. The curve was generated using the same gBlock artificial double-stranded DNA construct from Roux et al. (2020). Serial dilutions ranging from 0.001 to 1000 pg/L were amplified in triplicate, pooled and added to each MiSeq run. Technical replicates were produced with the same molecular identification tag and were not analyzed separately. The standard curves were analyzed in R (R Core Team, 2018) using linear regression of log-transformed raw read numbers and DNA concentrations (lm in *stats*). The limit of detection and limit of quantification could not calculated for tNGS in the same way as for qPCR assays because multiple runs were not performed on each sample. However, a minimum number of sequence copies in a sample, below which sequences are discarded, must be set to remove PCR artifacts and potential contamination. We ensured that chimeric sequences would not persist in the data after demultiplexing by using each primer only once. Knowing that the maximum number of green crab reads returned in blanks was one, the minimum number of raw reads was set at five for a positive detection. This number was also set as the minimum limit for the standard curves to calculate eDNA concentration. This threshold will remove error due to contamination quantified by negative controls, yet is still low enough to detect the presence of species’ with very low starting eDNA amounts.

### 2.4 Multi-scale eDNA occupancy modelling

Bayesian MCMC estimation of multi-scale eDNA occupancy modelling parameters was performed for tNGS and qPCR using the R package eDNAoccupancy (Dorazio & Erickson, 2018). The hierarchical parameters estimated by the models were: (1) the probability of green crab at a site (ψ); (2) the conditional probability of green crab in a sample given it is present at a site (θ); and (3) the conditional probability of green crab in a replicate of a sample given that it is present in the sample (ρ). The 6 site replicates per day of the *replicate* data set were pooled into 12 samples for each of 4 sites. Presence was determined by a returned C_t_ value for at least one qPCR replicate and number of reads >100 for tNGS. Two models were fit to each data set: the first model assumes that all parameters are constant and the second model assumes that ψ and ρ are constant whilst assuming that θ varies as a function of green crab abundance (CPUE). Models were run for 33,000 (tNGS models) and 11,000 (qPCR models) MCMC iterations. Convergence was achieved when the chain mixed well (“hairy” trace plot) and autocorrelation reached zero after discarding 10% burnin iterations. The posterior predictive loss criteria (Gelfand & Ghosh, 1998) (Gelfand and Ghosh 1998) and widely applicable information criteria (Watanabe, 2010, 2013) (Watanabe 2010, 2013) were used to assess if adding the CPUE covariate improved model fit without significantly increasing the variance, the smallest values indicating best fit. Posterior summaries of hierarchical occupancy parameters were calculated after discarding 10% burnin iterations.

### 2.5 Spatial patterns of CPUE and eDNA concentration

Generalized linear models (GLMs) were used to assess if spatial patterns of eDNA concentration derived from tNGS and qPCR assays correlated with abundance from traditional trapping surveys. The eDNA sampling regime included non-independent sample site replicates. Therefore, we also generated generalized linear mixed effect models (GLMM) with a random parameter to allow estimation of variance within and among groups of sample site replicates. We then selected the best fit model, GLM or GLMM, using the Hausman Specification Test (ph-test in plm) (Croissant & Millo, 2017) modified for output from lme4 (code available upon request). Residual plots, QQ-plots, and the Shapiro-Wilks test were used to assess the model fit between ordinary least squares log-transformed eDNA concentrations, Gaussian log-linked, and Gamma log-linked distributions for GLMs and GLMMs. *Site* and *replicate* data sets were analyzed separately for tNGS and qPCR assays, and only GLMs were used for *site* data sets because there was no site replication.

### 2.6 Partitioning sources of variation

It is important to determine the extent to which the spatial and temporal scales used during sampling contribute to variation in eDNA concentration. At the lowest sampling level, we expect that variation in qPCR technical replicates (three qPCR replicates of each site replicate) to be very small (not comparable with tNGS where technical replicates were assigned the same molecular identification tagged and pooled). We expect that variation among site replicates within each day will be greater than among technical replicates but smaller than variance among sites or between days. Variation between days is also expected to be less than among sites as eDNA concentration is not expected to change appreciably over a 24 hour period. The between-site variation was calculated to understand the total variation across Barkley Sound.

### 2.7 Abundance classification using eDNA concentration

One main goal of a quantitative eDNA assay is to estimate the relative abundance of a species from a set of observed eDNA concentrations. An informative approach would be to classify unknown sites into categories of relative abundance by using a panel of controls that include eDNA from multiple sites of known abundance. An ANOVA was used to test if log transformed eDNA concentrations from high and low abundance sites were significantly different. Mayne Bay and Pipestem Inlet were pooled into the high abundance category and Ritherdon Bay and San Mateo Bay were pooled into the low abundance category. Classification of sites was performed using linear discriminant analysis (lda in MASS) (R Core Team, 2018) (R Core Team) on log transformed eDNA concentrations from the tNGS *replicate* data set. Mayne Bay (high abundance) and Ritherdon Bay (low abundance) sites were used as training data to assess the posterior probabilities of classifying samples from Pipestem Inlet and San Mateo Bay into their correct categories of high and low abundance, respectively.

## 3.0 Results

### 3.1 Run statistics and standard curves

Two MiSeq runs resulted in ∼28.2 million raw paired reads, reduced to ∼25.6 million reads after removing low quality sequences, aligning paired ends, demultiplexing, and removing non-green crab or gBlock sequences. The proportion of PhiX reads without an adapter was as expected at about 15% across runs, however, almost all the reads that had a sample adapter and were not green crab or gBlock (∼1.9 million) appeared to be bleed-through PhiX from the spike-in. This is known to occur where a non-indexed PhiX is situated next to an indexed cluster on the flow cell (Hussman, 2015) and the high percentage spike-in created a large proportion of bleed-through in this case. For future runs, it is likely possible to reduce the spike-in to the recommended <5% and rely on other positional diversity elements built into the adapters and pairing scheme (Naik et al., 2020).

Standard curves on two tNGS MiSeq runs indicated 96.2% and 95.6% of the total variance was explained by a linear regression model of number of reads and log transformed gBlock PCR template concentration. The lowest concentrations for both curves of 0.002 copies/µL were removed because MiSeq Run #1 returned a single read, therefore the lowest concentration used in the standard curve for calculating eDNA concentration from reads was 0.02 copies/µL and 0.18 copies per reaction (three pooled PCR replicates). The LODs for the qPCR assay reported in Roux et al. (2020) were 2.01 copies/ µL (C_t_=35 +- 0.97) for 100% of replicates and 0.2 copies/µL (C_t_ = 38 +- 0.61) for 50% of replicates (Biggs et al., 2015; Polinski et al., 2015). Since the two methods cannot be directly compared, we report that the tNGS method detected the green crab gBlock across two MiSeq runs at a concentration that was 10-100 times lower than the limit of quantification and the limit of detection for qPCR.

### 3.2 Catch per unit effort and eDNA concentrations from qPCR and tNGS for samples and blanks

Catch per unit effort (CPUE) was 2.0 and 7.4 at low abundance sites (San Mateo Bay and Ritherdon Bay, respectively) and 23.0, 35.4, and 73.5 at high abundance sites (Mayne Bay, Pipestem Inlet, and Hillier Island, respectively) (Fig. 1). Zooplankton samples returned all non-detects for qPCR and almost zero reads for tNGS so this sample type was removed from further analyses (data not shown). eDNA concentrations of sample site replicates and across days at each site clustered tightly together for the low abundance sites for the tNGS assay, but the high abundance sites did not cluster as tightly (*replicate* and *site* data, Fig. 2a and 2c, respectively). Despite the variation in eDNA concentrations measured at high abundance sites, there was distinct separation between low and high abundance sites for the tNGS assay but not for the qPCR assay, where quartiles significantly overlapped among most sites (Fig. 2b and 2d). All blank samples returned one or zero reads for green crab.

**Fig. 2.**
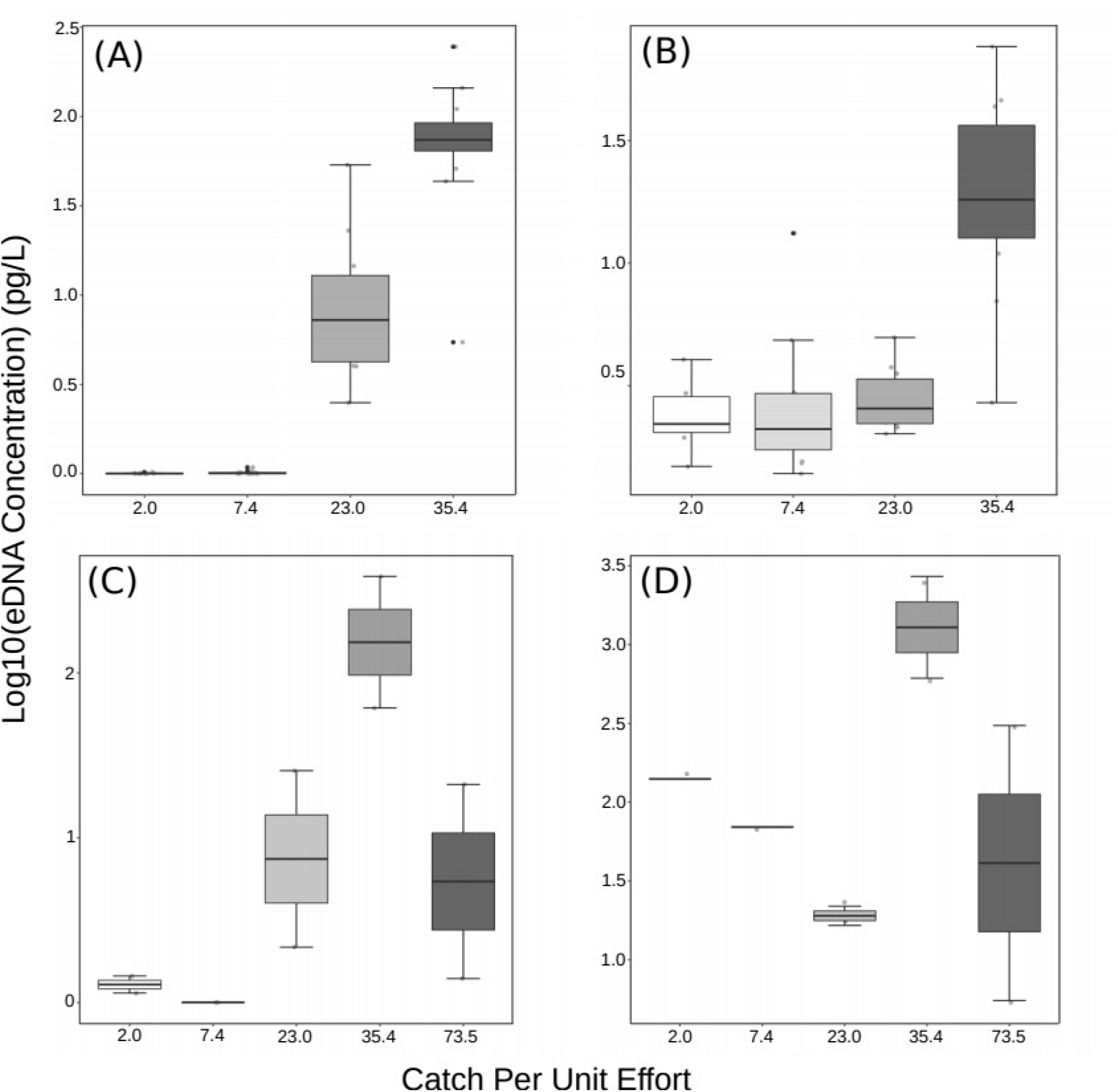
Box and dotplots of eDNA concentration by site marked by catch per unit effort (CPUE). *Replicate* tNGS (a) and qPCR (b) data set. *Site* tNGS (c) and qPCR (d) data set. CPUE for low abundance sites are: 2.0 = San Mateo Bay and 7.4 = Ritherdon Bay. CPUE for high abundance sites are: 23.0 = Mayne Bay, 35.4 = Pipestem Inlet, and 73.5 = Hilliers Island.

### 3.3 Multi-scale eDNA occupancy modelling

Model selection criteria for tNGS and qPCR indicated that allowing the detection probability of eDNA in a sample to vary with green crab relative abundance (CPUE) improved model fit (Supplementary Table S3). CPUE and the conditional probability of eDNA detection in a sample (θ) were positively correlated for both methods (median alpha.cpue ± SE: tNGS = 0.799 ± 0.007, qPCR = 0.857 ± 0.012). The probabilities of detecting green crab at the high CPUE sites did not differ appreciably between methods (Mayne Bay: median [95% CI] = 0.950 [0.832,0.997] qPCR, 0.972 [0.872,0.999] tNGS, Pipestem Inlet: median [95% CI] = 0.999 [0.882,1.000] qPCR, 0.993 [0.901,1.000] tNGS) (Fig. 3). However, tNGS had higher probabilities of detecting green crab at the low abundance sites (San Mateo Bay: median [95% CI] = 0.699 [0.467,0.883] qPCR, 0.798 [0.553,0.994] tNGS, Ritherdon Bay: median [95% CI] = 0.795 [0.627,0.918] qPCR, 0.868 [0.709,0.994] tNGS) (Fig. 3).

**Fig. 3.**
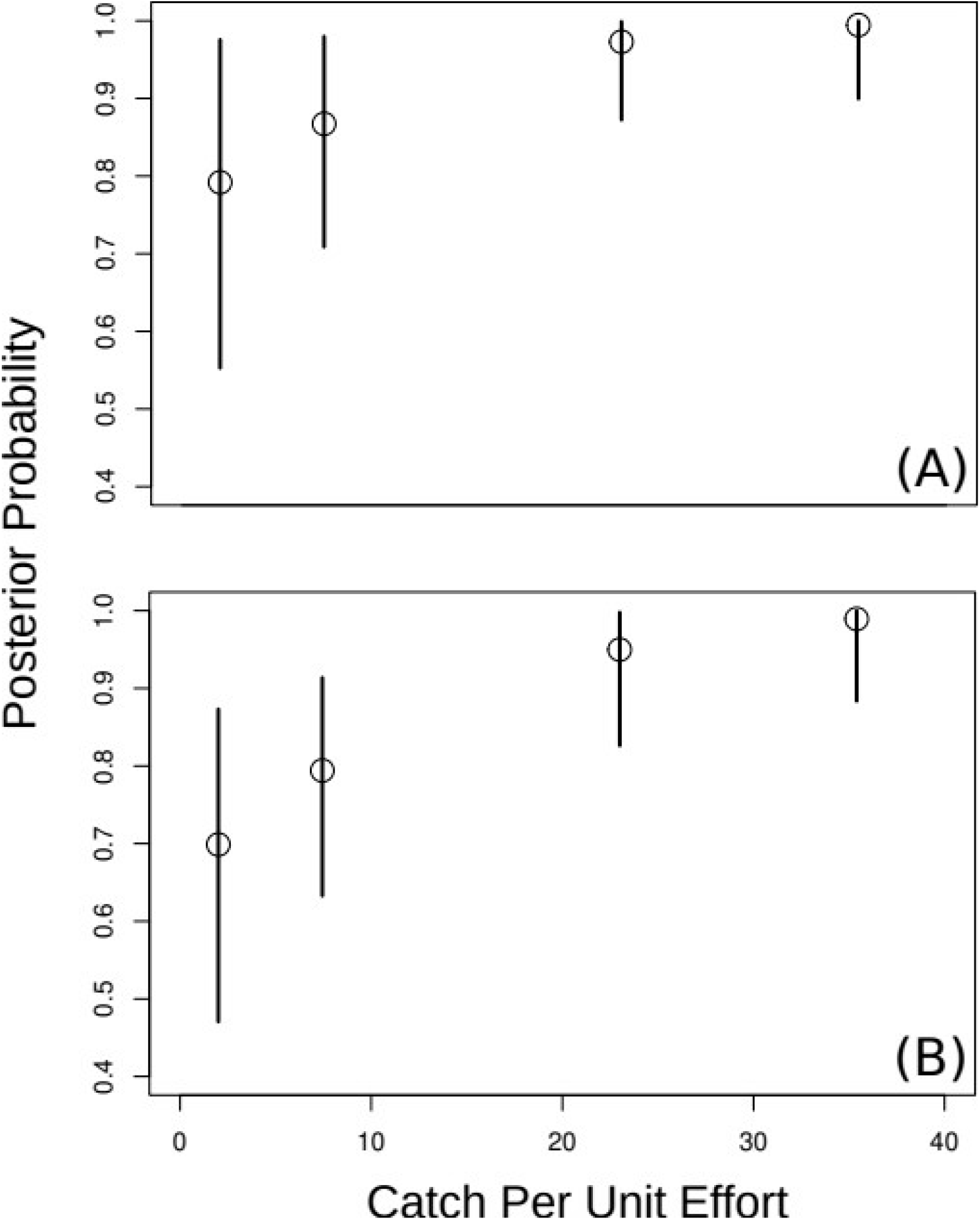
Estimated probabilities of occurrence of green crab eDNA at sites with different catch per unit effort (CPUE) values. Symbols are estimates of posterior medians with 95% credible intervals. Models with CPUE covariates for tNGS (a) and qPCR (b).

### 3.4 Spatial patterns of CPUE and eDNA concentration

*Site* and *replicate* data sets for qPCR and tNGS assays were all best fit by an OLS linear regression model of log transformed eDNA concentrations against CPUE. CPUE was significantly correlated with eDNA concentration for both assays; however, the tNGS data was more precise and had less error (Adjusted R^2^ = 0.86) relative to the qPCR assay (Adjusted R^2^ = 0.48) (Table 1). Predicted CPUE from tNGS eDNA concentration was more accurate relative to qPCR with much smaller 95% confidence intervals, especially at low abundance sites (Supplementary Fig. S1). CPUE was not correlated with eDNA concentration for either assay using the *site* data (Table 1).

**Table 1.** Linear regression results for tNGS and qPCR eDNA concentration against CPUE performed on *site* and *replicate* data sets.

| *Predictors* | *tNGS (replicate)* |  | *qPCR (replicate)* |  | *tNGS (site)* |  | *qPCR (site)* |  |
| --- | --- | --- | --- | --- | --- | --- | --- | --- |
|  | *Estimates* | *CI* | *Estimates* | *CI* | *Estimates* | *CI* | *Estimates* | *CI* |
| (Intercept) | -0.27*** | -0.41 –<br>-0.13 | 0.19* | 0.01 –<br>0.37 | 0.42 | -0.42 –<br>1.26 | 2.08** | 0.96 –<br>3.20 |
| CPUE | 0.06*** | 0.05 –<br>0.06 | 0.03*** | 0.02 –<br>0.03 | 0.01 | -0.01 –<br>0.03 | -0.00 | -0.03 –<br>0.02 |
| Observations | 48 |  | 41 |  | 10 |  | 8 |  |
| R<sup>2</sup> | 0.867 |  | 0.493 |  | 0.137 |  | 0.005 |  |
| Adjusted R<sup>2</sup> | 0.864 |  | 0.480 |  | 0.029 |  | -0.161 |  |

CI is 95% confidence interval. Significance \*\*\* = $p < 0.001$ , \*\* = $p < 0.05$ , \* = $p < 0.1$ .

### 3.5 Sources of variation

Total within-site variation for tNGS (site replicates) was less than qPCR (combined technical qPCR replicates and site replicates) (mean SD [90%CI] = tNGS: 0.07 [0.06,0.09]; qPCR: 0.10 [0.08,0.11]), respectively). For both methods, mean within-site variation was only slightly less than the variation between days at each site (mean SD [90%CI] = qPCR = 0.11 [0.09,0.13], tNGS = 0.09 [0.07,0.10]), and was much less than the variation among sites for each day (mean SD [90%CI] = qPCR = 0.19 [0.16,0.23], tNGS = 0.35 [0.30,0.41]) (Fig. 4). It is important that the variation between days was similar to variation between site replicates, showing that eDNA concentration did not change appreciably over a tidal cycle. The difference between within-site variation and between-site variation was greater for tNGS than qPCR, suggesting that tNGS has more power to detect differences among sites relative to qPCR.

**Fig. 4.**
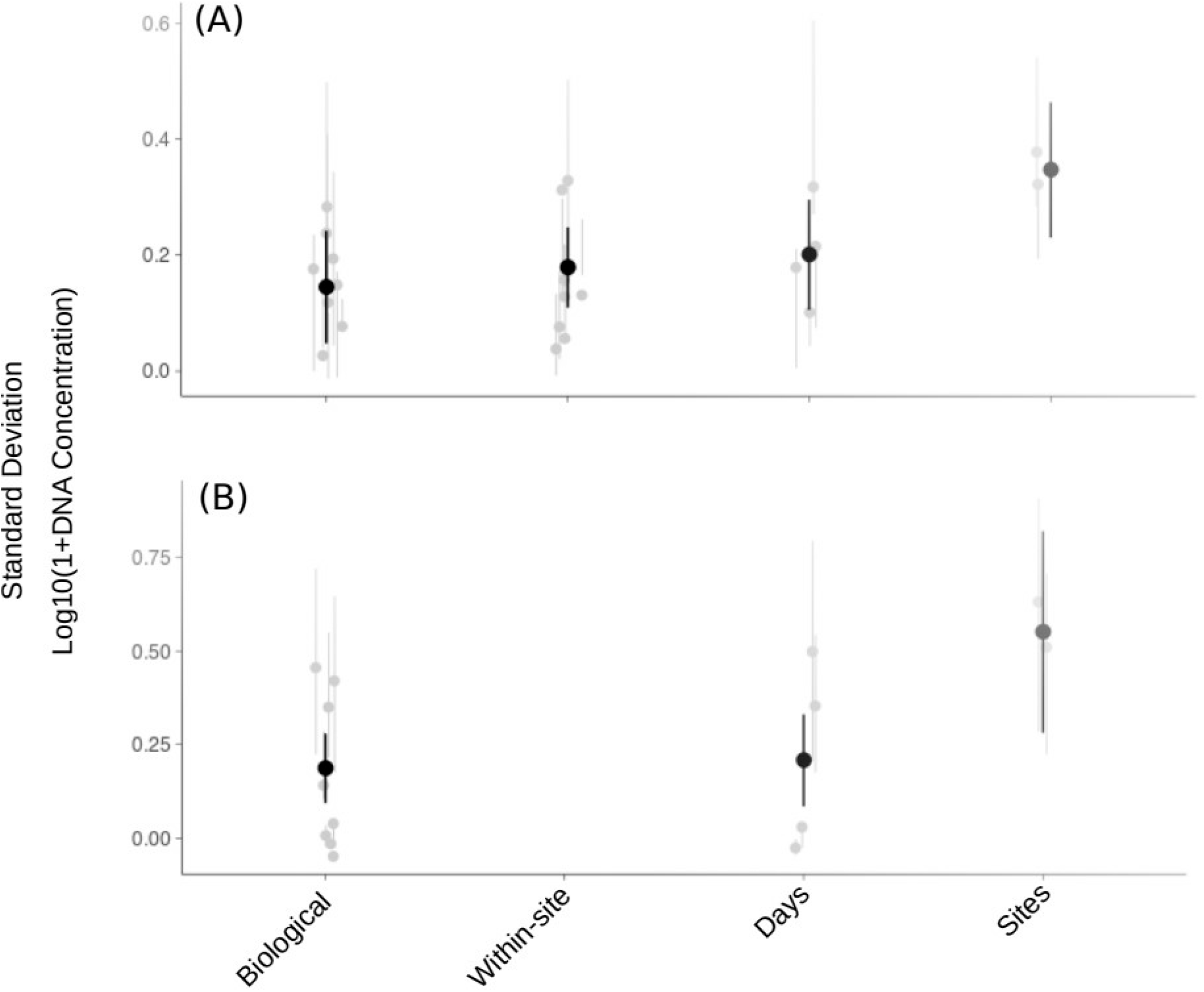
Partitioning sources of variation for qPCR (a) and tNGS (b) green crab eDNA surveys, the *replicate* data set only. Grey points show estimated values in each category and black points show within category means ± 90% confidence interval. For both assays ‘Biological’ replication is six site replicates per site per day. Within-site (‘W-S’) variation for qPCR is the addition of ‘Biological’ and ‘Technical’ (qPCR technical replicates) and represents all variance within sites. For tNGS the total within-site variation is the ‘Biological’ variation. Variation between days within each site is ‘Days’, and the among site variation in eDNA concentration for each day is ‘Sites’.

### 3.6 Abundance classification using eDNA concentration

The ANOVA revealed a significant difference in mean eDNA concentrations between high and low abundance sites compared to variation within categories (F_1_ = 219.2, P < 0.001). Linear discriminant analysis was highly successful at classifying samples to high and low abundance classes with all samples except one being correctly classified with a posterior probability of 99.9% or higher (one sample had a posterior probability of 95.8%). Thus relative abundance was correctly classified despite variation observed in eDNA concentration across days and site replicates.

## 4.0 Discussion

Improving species detection rates and quantification precision from eDNA substantially augments the benefits molecular surveys offer management programs seeking alternatives to labour intensive traditional survey techniques. For rare or scarce species, molecular traces are often the only available information on which to base management decisions and corresponding actions. Integrating eDNA into the management toolbox requires a deep understanding of how eDNA behaves in space and time and with respect to traditional metrics. The current study has used the invasive European green crab in the northeast Pacific to showcase the utility of eDNA as a tool for determining species presence and estimating relative abundance, even at very low population densities. Here we described the new tNGS method that outperformed the widely used qPCR method for detecting green crab at low densities, correlating with data from traditional trapping methods, and reflecting the differential abundances of crabs across sample sites in Barkley Sound. Here we discuss each of these new advances in detail and provide practical advice on how best to apply this new assay to specific management objectives.

### 4.1 Standard curves and limit of detection

The proof of concept of direct high throughput sequencing of PCR amplicons without normalization was validated by repeatable quantification; two standardized curves of DNA concentration against sequencing read numbers from two Illumina platform runs had R2 values close to 1 (Mauvisseau, Burian, et al., 2019). Reads were returned for gBlock controls at 0.02 copies/µL (0.18 copies per reaction) for tNGS, which was 10 times lower than the LOD (>50% of replicates positive) of the green crab qPCR assay of Roux et al. (2020). Reported LODs of highly sensitive qPCR assays vary considerably but >0.5 copies/µL has been reported as a minimum for positive detection in some studies (Baker et al., 2018; Mauvisseau, Davy-Bowker, et al., 2019). The tNGS assay could therefore exceed the sensitivity of both qPCR and dPCR whilst eliminating false positives from background fluorescence and non-specific amplification. It also provides more information on the target detected because a sequence is generated, allowing for population level analyses.

### 4.2 Occupancy modelling in tNGS and qPCR assays

Estimates of green crab occurrence and detection probabilities were fairly robust for both assays at high eDNA concentrations; Mayne Bay and Pipestem Inlet had median probabilities between 95% and 99% with very little variation among site replicates. At low abundances (San Mateo Bay and Ritherdon Bay), the tNGS assay showed consistently higher detection probabilities with a median of 7%-10% greater than qPCR. Green crab CPUE as a measure of relative abundance was identified as an important covariate for explaining θ_eDNA_, inferring that species detection increases as species abundance increases. This relationship has been observed in some (Schmelzle & Kinziger, 2016) but not all eDNA studies (Goldberg et al., 2011). This result showcases some of the problems of fluorescence based quantification at low template concentrations and suggests that tNGS could be more accurate in cases where eDNA availability is expected to be low.

### 4.3 Spatial distribution of eDNA concentration and relationships with catch data

There is a clear difference in the pattern of eDNA concentrations between assays across sites and CPUE values; tNGS tends to reflect increasing CPUE whereas qPCR seems to reach a minimum abundance level before exhibiting an increase in eDNA concentration. The linear relationships break down when site replicates are pooled, enforcing the need for site replication to reduce stochastic effects in eDNA (Cannon et al., 2016) and PCR (Kebschull & Zador, 2015).

The significant positive relationships observed between eDNA concentration and CPUE for qPCR and tNGS assays support the use of eDNA as a metric for species abundance, an important result that has been established in multiple previous studies and confirmed here for green crab (Doi et al., 2017; Knudsen et al., 2019; Lacoursière-Roussel et al., 2016; Lehman et al., 2019; Shelton et al., 2019). The tNGS method shows higher precision relative to qPCR for predicting CPUE, and this difference is most apparent at low abundances where the predicted abundance (CPUE) confidence interval for tNGS is much smaller than qPCR.

### 4.4 Partitioning variance in tNGS and qPCR assays

The variance partitions showed expected patterns for both assays, with mean total within-site variation being the smallest values and increasing only slightly between days. This supports previous evidence that eDNA availability is endogenous to each site after undergoing diurnal tidal cycles (Kelly et al., 2018), with just marginally higher variance over a 24 hour period compared to total within-site variation in each day. The jump in between-site variation for both assays was expected based on the large differences in abundance levels among sites specifically targeted with the sampling design. The tNGS assay showed a greater difference between within-site and between-site variation relative to qPCR, suggesting that tNGS has greater power to detect spatial variation in eDNA concentration. In applications where abundance may not be known a priori or estimated with another method, the variance between sites may be smaller.

### 4.5 Field sampling considerations

Comparisons between *site* and *replicate* data sets indicate that increasing the number of replicates at each sample site increases detection probability and predictive capacity for relative abundance, and the effects are much more significant at low species abundance. Sampling across days did not affect the estimated eDNA concentration greatly. This is consistent with results from other studies (Kelly et al., 2018), hence it does not seem necessary to capture small-scale temporal variation (i.e. days) to accurately estimate abundance. However, seasonal or inter-annual variability will likely be an important consideration for some sampling designs depending on research objectives and the target organisms.

Community DNA from zooplankton tows contained very little to no detectable green crab DNA, contrary to previous studies that show Arthropod diversity is higher in DNA samples from zooplankton versus water (Koziol et al., 2018; Westfall et al., 2019). This indicates the eDNA signal detected here was largely from adults and that larval seasonality, while becoming important during some periods, did not affect the results of this survey. It is helpful for integration into management programs to ensure that the eDNA signal measured is largely independent of seasonally fluctuating larval densities.

### 4.6 Conclusions and applying eDNA to current management programs

Managing green crab in the northeast Pacific involves ongoing monitoring of population abundance at invaded sites as a census or in response to eradication efforts (Duncombe & Therriault, 2017), and early detection of secondary spread into new environments (Roux et al. 2020). An eDNA-based qPCR assay was developed by Roux et al. (2020) in part to meet these management needs; however, the tNGS assay presented here has demonstrated increased performance for detecting green crab at low abundance, detecting changes in abundance among sites, and predicting abundance as measured by traditional catch data. Standard DNA concentrations that were 10-to 100-times lower than the qPCR limit of detection returned significant numbers of sequencing reads in the tNGS assay, and in our field assessments this translated to an increase of 7% to 10% detection probability of green crab at sites with low abundance. Although this number seems marginal, even a small increase in detection probability could make a consequential impact at the first onset of potentially damaging secondary spread (Dougherty et al., 2016; Holman et al., 2019). Further, although these trade-offs will vary among monitoring programs and labs due to unique particularities and needs therein, we expect tNGS could bring some operational advantages with respect to time- and cost-efficiencies compared to qPCR for programs processing high sample numbers, where high sensitivity and/or assay specificity are important considerations.

Using the tNGS assay to quantify relative species abundance is a major step forward for management efforts of European green crab and other invaders. Implementing the assay for this purpose could be achieved with linear discriminant analysis as was successfully demonstrated here using a subset of sites to train the classifier. Including a panel of internal controls that represent most of the expected variance enables the classification of sites with no *a priori* information into categories of relative abundance. Not only can this approach be used to assess presence and abundance as part of ongoing monitoring of established or nascent populations, it can also be used to assess the efficacy of eradication efforts when other methods may suggest the job is done.

Continuing to validate the tNGS assay with such promising results should be the focus of future research. There have been recent calls for standardizing determination of the limit of detection for eDNA-based qPCR assays (Roux et al., 2020), especially at low concentrations where MIQE guidelines are restrictive (Hunter et al., 2018). Future research using tNGS could involve setting a standard guideline for quantifying the limit of detection by assessing behaviour across multiple assays – information that could be conveyed to managers to better guide decision making.

## Acknowledgements

The authors thank Dr. Kristina Miller-Saunders and Angela Schulze for MiSeq assistance, and Kara Aschenbrenner, Geoff Lowe and Louise Roux for field and laboratory assistance. Funding was provided to Drs. Abbott and Therriault from Fisheries and Oceans Canada’s Program for Aquaculture Regulatory Research (project no. PARR-2016-P-03) and funding to Dr. Therriault from Fisheries and Oceans Canada’s Aquatic Invasive Species Monitoring program in the Pacific Region.

## References

Alam, M. M. M., Westfall, K. M., & Pálsson, S. (2017). Historical demography and genetic differentiation of the giant freshwater prawn Macrobrachium rosenbergii in Bangladesh based on mitochondrial and ddRAD sequence variation. Ecology and Evolution, 7(12), 4326–4335. https://doi.org/10.1002/ece3.3023

Baker, C. S., Steel, D., Nieukirk, S., & Klinck, H. (2018). Environmental DNA (eDNA) From the Wake of the Whales: Droplet Digital PCR for Detection and Species Identification. Frontiers in Marine Science. https://doi.org/10.3389/fmars.2018.00133

Biggs, J., Ewald, N., Valentini, A., Gaboriaud, C., Dejean, T., Griffiths, R. A., Foster, J., Wilkinson, J. W., Arnell, A., Brotherton, P., Williams, P., & Dunn, F. (2015). Using eDNA to develop a national citizen science-based monitoring programme for the great crested newt (Triturus cristatus). Biological Conservation, 183, 19–28. https://doi.org/10.1016/j.biocon.2014.11.029

Boyer, F., Mercier, C., Bonin, A., Le Bras, Y., Taberlet, P., & Coissac, E. (2016). obitools: a unix-inspired software package for DNA metabarcoding. Molecular Ecology Resources, 16(1), 176– 182. https://doi.org/10.1111/1755-0998.12428

Bustin, S. A., Benes, V., Garson, J. A., Hellemans, J., Huggett, J., Kubista, M., Mueller, R., Nolan, T., Pfaffl, M. W., Shipley, G. L., Vandesompele, J., & Wittwer, C. T. (2019). The MIQE Guidelines: Minimum Information for Publication of Quantitative Real-Time PCR Experiments. Clinical Chemistry, 55(4), 611–622. https://doi.org/10.1373/clinchem.2008.112797

Buxton, A. S., Groombridge, J. J., Zakaria, N. B., & Griffiths, R. A. (2017). Seasonal variation in environmental DNA in relation to population size and environmental factors. Scientific Reports, 7, 46294. https://doi.org/10.1038/srep46294

Camacho, C., Coulouris, G., Avagyan, V., Ma, N., Papadopoulos, J., Bealer, K., & Madden, T. L. (2009). BLAST+: architecture and applications. BMC Bioinformatics, 10, 421. https://doi.org/10.1186/1471-2105-10-421

Cannon, M. V, Hester, J., Shalkhauser, A., Chan, E. R., Logue, K., Small, S. T., & Serre, D. (2016). In silico assessment of primers for eDNA studies using PrimerTree and application to characterize the biodiversity surrounding the Cuyahoga River. Scientific Reports, 6(1), 22908. https://doi.org/10.1038/srep22908

Carraro, L., Hartikainen, H., Jokela, J., Bertuzzo, E., & Rinaldo, A. (2018). Estimating species distribution and abundance in river networks using environmental DNA. Proceedings of the National Academy of Sciences, 115(46), 11724 LP – 11729. https://doi.org/10.1073/pnas.1813843115

Clement, M. J., O’Keefe, J. M., & Walters, B. (2015). A method for estimating abundance of mobile populations using telemetry and counts of unmarked animals. Ecosphere, 6(10), art184. https://doi.org/10.1890/ES15-00180.1

Croissant, Y., & Millo, G. (2017). Panel Data Econometrics in R: The plm Package. Journal of Statistical Software, 27(2), 1–43. https://doi.org/0.18637/jss.v027.i02

Davis, A., Hooten, M., Miller, R., Farnsworth, M., Lewis, J., Moxcey, M., & Pepin, K. (2016). Inferring invasive species abundance using removal data from management actions. Ecological Applications, 26(7), 2339–2346. https://doi.org/10.1002/eap.1383

Deiner, K., Bik, H. M., Mächler, E., Seymour, M., Lacoursière-Roussel, A., Altermatt, F., Creer, S., Bista, I., Lodge, D. M., de Vere, N., Pfrender, M. E., & Bernatchez, L. (2017). Environmental DNA metabarcoding: Transforming how we survey animal and plant communities. Molecular Ecology, 26(21), 5872–5895. https://doi.org/10.1111/mec.14350

Doi, H., Inui, R., Akamatsu, Y., Kanno, K., Yamanaka, H., Takahara, T., & Minamoto, T. (2017). Environmental DNA analysis for estimating the abundance and biomass of stream fish. Freshwater Biology, 62(1), 30–39. https://doi.org/10.1111/fwb.12846

Dorazio, R. M., & Erickson, R. A. (2018). ednaoccupancy: An r package for multiscale occupancy modelling of environmental DNA data. Molecular Ecology Resources, 18(2), 368–380. https://doi.org/10.1111/1755-0998.12735

Dougherty, M. M., Larson, E. R., Renshaw, M. A., Gantz, C. A., Egan, S. P., Erickson, D. M., & Lodge, D. M. (2016). Environmental DNA (eDNA) detects the invasive rusty crayfish Orconectes rusticus at low abundances. Journal of Applied Ecology, 53(3), 722–732. https://doi.org/10.1111/1365-2664.12621

Duncombe, L. G., & Therriault, T. W. (2017). Evaluating trapping as a method to control the European green crab, Carcinus maenas, population at Pipestem Inlet, British Columbia. Management of Biological Invasions, 8(2), 235–246. https://doi.org/10.3391/mbi.2017.8.2.11

Elbrecht, V., & Leese, F. (2015). Can DNA-Based Ecosystem Assessments Quantify Species Abundance? Testing Primer Bias and Biomass—Sequence Relationships with an Innovative Metabarcoding Protocol. PLOS ONE, 10(7), e0130324. https://doi.org/10.1371/journal.pone.0130324

Ellison, S. L. R., English, C. A., Burns, M. J., & Keer, J. T. (2006). Routes to improving the reliability of low level DNA analysis using real-time PCR. BMC Biotechnology, 6, 33. https://doi.org/10.1186/1472-6750-6-33

Forsström, T., & Vasemägi, A. (2016). Can environmental DNA (eDNA) be used for detection and monitoring of introduced crab species in the Baltic Sea? Marine Pollution Bulletin, 109(1), 350– 355. https://doi.org/10.1016/j.marpolbul.2016.05.054

Franklin, T. W., McKelvey, K. S., Golding, J. D., Mason, D. H., Dysthe, J. C., Pilgrim, K. L., Squires, J. R., Aubry, K. B., Long, R. A., Greaves, S. E., Raley, C. M., Jackson, S., MacKay, P., Lisbon, J., Sauder, J. D., Pruss, M. T., Heffington, D., & Schwartz, M. K. (2019). Using environmental DNA methods to improve winter surveys for rare carnivores: DNA from snow and improved noninvasive techniques. Biological Conservation, 229, 50–58. https://doi.org/10.1016/j.biocon.2018.11.006

Furlan, E. M., Gleeson, D., Wisniewski, C., Yick, J., & Duncan, R. P. (2019). eDNA surveys to detect species at very low densities: A case study of European carp eradication in Tasmania, Australia. Journal of Applied Ecology, 56(11), 2505–2517. https://doi.org/10.1111/1365-2664.13485

Gelfand, A. E., & Ghosh, S. K. (1998). Model Choice: A Minimum Posterior Predictive Loss Approach. Biometrika, 85(1), 1–11. http://www.jstor.org/stable/2337305

Gillespie, G. E., Phillips, A. C., Paltzat, D. L., & Therriault, T. W. (2007). Status of the European green crab, Carcinus maenas, in British Columbia - 2006.

Goldberg, C. S., Pilliod, D. S., Arkle, R. S., & Waits, L. P. (2011). Molecular Detection of Vertebrates in Stream Water: A Demonstration Using Rocky Mountain Tailed Frogs and Idaho Giant Salamanders. PLOS ONE, 6(7), e22746. https://doi.org/10.1371/journal.pone.0022746

Haines, A. M., Zak, M., Hammond, K., Scott, J. M., Goble, D. D., & Rachlow, J. L. (2013). Uncertainty in Population Estimates for Endangered Animals and Improving the Recovery Process. Animals: An Open Access Journal from MDPI, 3(3), 745–753. https://doi.org/10.3390/ani3030745

Harper, L. R., Lawson Handley, L., Hahn, C., Boonham, N., Rees, H. C., Gough, K. C., Lewis, E., Adams, I. P., Brotherton, P., Phillips, S., & Hänfling, B. (2018). Needle in a haystack? A comparison of eDNA metabarcoding and targeted qPCR for detection of the great crested newt (Triturus cristatus). Ecology and Evolution, 8(12), 6330–6341. https://doi.org/10.1002/ece3.4013

Harvey, C. T., Qureshi, S. A., & MacIsaac, H. J. (2009). Detection of a colonizing, aquatic, non-indigenous species. Diversity and Distributions, 15(3), 429–437. https://doi.org/10.1111/j.1472-4642.2008.00550.x

Holman, L. E., de Bruyn, M., Creer, S., Carvalho, G., Robidart, J., & Rius, M. (2019). Detection of introduced and resident marine species using environmental DNA metabarcoding of sediment and water. Scientific Reports, 9(1), 11559. https://doi.org/10.1038/s41598-019-47899-7

Howald, G., Donlan, C. J., Galvan, J. P., Russell, J. C., Parkes, J., Samaniego, A., Wang, Y., Veitch, D., Genovesi, P., Pascal, M., Saunders, A., & Tershy, B. (2007). Invasive Rodent Eradication on Islands. Conservation Biology, 21(5), 1258–1268. https://doi.org/10.1111/j.1523-1739.2007.00755.x

Hunter, M., Meigs-Friend, G., Ja, F., A, T. K., Rm, D., Keith-Diagne, L., Luna, F., Jm, L., & P, R. J. (2018). Surveys of environmental DNA (eDNA): a new approach to estimate occurrence in Vulnerable manatee populations. Endangered Species Research, 35, 101–111. https://www.int-res.com/abstracts/esr/v35/p101-111/

Hussman, J. (2015). Expanding the applications of high-throughput DNA sequencing [University of Texas at Austin]. http://hdl.handle.net/2152/31375

Jones, J. P. G. (2011). Monitoring species abundance and distribution at the landscape scale. Journal of Applied Ecology, 48, 9–13.

Kebschull, J. M., & Zador, A. M. (2015). Sources of PCR-induced distortions in high-throughput sequencing data sets. Nucleic Acids Research, 43(21), e143–e143. https://doi.org/10.1093/nar/gkv717

Kelly, R. P., Gallego, R., & Jacobs-Palmer, E. (2018). The effect of tides on nearshore environmental DNA. PeerJ, 6, e4521–e4521. https://doi.org/10.7717/peerj.4521

Klymus, K. E., Merkes, C. M., Allison, M. J., Goldberg, C. S., Helbing, C. C., Hunter, M. E., Jackson, C. A., Lance, R. F., Mangan, A. M., Monroe, E. M., Piaggio, A. J., Stokdyk, J. P., Wilson, C. C., & Richter, C. A. (2020). Reporting the limits of detection and quantification for environmental DNA assays. Environmental DNA, 2(3), 271–282. https://doi.org/10.1002/edn3.29

Knudsen, S. W., Ebert, R. B., Hesselsøe, M., Kuntke, F., Hassingboe, J., Mortensen, P. B., Thomsen, P. F., Sigsgaard, E. E., Hansen, B. K., Nielsen, E. E., & Møller, P. R. (2019). Species-specific detection and quantification of environmental DNA from marine fishes in the Baltic Sea. Journal of Experimental Marine Biology and Ecology, 510, 31–45. https://doi.org/10.1016/j.jembe.2018.09.004

Koziol, A., Stat, M., Simpson, T., Jarman, S., DiBattista, J. D., Harvey, E. S., Marnane, M., McDonald, J., & Bunce, M. (2018). Environmental DNA metabarcoding studies are critically affected by substrate selection. Molecular Ecology Resources, 0(0). https://doi.org/10.1111/1755-0998.12971

Krebs, C. J. (1972). Ecology: the experimental analysis of distribution and abundance / Charles J. Krebs. Harper & Row.

Lacoursière-Roussel, A., Howland, K., Normandeau, E., Grey, E. K., Archambault, P., Deiner, K., Lodge, D. M., Hernandez, C., Leduc, N., & Bernatchez, L. (2018). eDNA metabarcoding as a new surveillance approach for coastal Arctic biodiversity. Ecology and Evolution, 8(16), 7763–7777. https://doi.org/10.1002/ece3.4213

Lacoursière-Roussel, A., Rosabal, M., & Bernatchez, L. (2016). Estimating fish abundance and biomass from eDNA concentrations: variability among capture methods and environmental conditions. Molecular Ecology Resources, 16(6), 1401–1414. https://doi.org/10.1111/1755-0998.12522

Langlois, V. S., Allison, M. J., Bergman, L. C., To, T. A., & Helbing, C. C. (2020). The need for robust qPCR-based eDNA detection assays in environmental monitoring and species inventories. Environmental DNA, n/a(n/a). https://doi.org/10.1002/edn3.164

Laramie, M. B., Pilliod, D. S., & Goldberg, C. S. (2015). Characterizing the distribution of an endangered salmonid using environmental DNA analysis. Biological Conservation, 183, 29–37. https://doi.org/10.1016/j.biocon.2014.11.025

Lehman, R. N., Poulakis, G. R., Scharer, R. M., Schweiss, K. E., Hendon, J. M., & Phillips, N. M. (2019). An environmental DNA tool for monitoring the status of the Critically Endangered Smalltooth Sawfish, <em>Pristis pectinata</em>, in the Western Atlantic. BioRxiv, 765321. https://doi.org/10.1101/765321

Lodge, D. M., Williams, S., MacIsaac, H. J., Hayes, K. R., Leung, B., Reichard, S., Mack, R. N., Moyle, P. B., Smith, M., Andow, D. A., Carlton, J. T., & McMichael, A. (2006). BIOLOGICAL INVASIONS: RECOMMENDATIONS FOR U.S. POLICY AND MANAGEMENT. Ecological Applications, 16(6), 2035–2054. https://doi.org/10.1890/1051-0761(2006)016[2035:BIRFUP]2.0.CO;2

Mauvisseau, Q., Burian, A., Gibson, C., Brys, R., Ramsey, A., & Sweet, M. (2019). Influence of accuracy, repeatability and detection probability in the reliability of species-specific eDNA based approaches. Scientific Reports, 9(1), 580. https://doi.org/10.1038/s41598-018-37001-y

Mauvisseau, Q., Davy-Bowker, J., Bulling, M., Brys, R., Neyrinck, S., Troth, C., & Sweet, M. (2019). Combining ddPCR and environmental DNA to improve detection capabilities of a critically endangered freshwater invertebrate. Scientific Reports, 9(1), 14064. https://doi.org/10.1038/s41598-019-50571-9

Naik, T., Sharda, M., & Pandit, A. (2020). High quality single amplicon sequencing method for illumina platforms using ‘N’ (0-10) spacer primer pool without PhiX spik-in. bioRxiv.

Pilliod D. S., Goldberg C. S., Arkle R. S., & Waits L. P. (2013). Estimating occupancy and abundance of stream amphibians using environmental DNA from filtered water samples. Canadian Journal of Fisheries and Aquatic Sciences, 70(8), 1123–1130. https://doi.org/10.1139/cjfas-2013-0047

Polinski, M., Lowe, G., Meyer, G., Corbeil, S., Colling, A., Caraguel, C., & Abbott, C. L. (2015). Molecular detection of Mikrocytos mackini in Pacific oysters using quantitative PCR. Molecular and Biochemical Parasitology, 200(1), 19–24. https://doi.org/10.1016/j.molbiopara.2015.04.004

R Core Team, R. (2018). R: A Language and Environment for Statistical Computing.

Roux, L.-M., Giblot-Ducray, D., Bott, M., Wiltshire, K., Deveney, M., Westfall, K., & Abbott, C. (2020). Analytical validation and initial field testing of qPCR assay for eDNA-based detection of invasive green crab (Carcinus maenas). In Environmental DNA.

Sassoubre, L. M., Yamahara, K. M., Gardner, L. D., Block, B. A., & Boehm, A. B. (2016). Quantification of Environmental DNA (eDNA) Shedding and Decay Rates for Three Marine Fish. Environmental Science & Technology, 50(19), 10456–10464. https://doi.org/10.1021/acs.est.6b03114

Schmelzle, M. C., & Kinziger, A. P. (2016). Using occupancy modelling to compare environmental DNA to traditional field methods for regional-scale monitoring of an endangered aquatic species. Molecular Ecology Resources, 16(4), 895–908. https://doi.org/10.1111/1755-0998.12501

Schubert, M., Lindgreen, S., & Orlando, L. (2016). AdapterRemoval v2: rapid adapter trimming, identification, and read merging. BMC Research Notes, 9(1), 88. https://doi.org/10.1186/s13104-016-1900-2

Shelton, A. O., Kelly, R. P., O’Donnell, J. L., Park, L., Schwenke, P., Greene, C., Henderson, R. A., & Beamer, E. M. (2019). Environmental DNA provides quantitative estimates of a threatened salmon species. Biological Conservation, 237, 383–391. https://doi.org/10.1016/j.biocon.2019.07.003

Strain, M. C., Lada, S. M., Luong, T., Rought, S. E., Gianella, S., Terry, V. H., Spina, C. A., Woelk, C. H., & Richman, D. D. (2013). Highly Precise Measurement of HIV DNA by Droplet Digital PCR. PLOS ONE, 8(4), e55943. https://doi.org/10.1371/journal.pone.0055943

Taberlet, P., Coissac, E., Hajibabaei, M., & Reiseberg, L. H. (2012). Environmental DNA. Molecular Ecology, 21(8), 1789–1793. https://doi.org/10.1111/j.1365-294X.2012.05542.x

Thomas, A. C., Tank, S., Nguyen, P. L., Ponce, J., Sinnesael, M., & Goldberg, C. S. (2019). A system for rapid eDNA detection of aquatic invasive species. Environmental DNA, n/a(n/a). https://doi.org/10.1002/edn3.25

Thomsen, P. F., Kielgast, J., Iversen, L. L., Møller, P. R., Rasmussen, M., & Willerslev, E. (2012). Detection of a Diverse Marine Fish Fauna Using Environmental DNA from Seawater Samples. PLOS ONE, 7(8), e41732. https://doi.org/10.1371/journal.pone.0041732

Thomsen, P. F., & Willerslev, E. (2015). Environmental DNA – An emerging tool in conservation for monitoring past and present biodiversity. Biological Conservation, 183, 4–18. https://doi.org/10.1016/j.biocon.2014.11.019

scales. Biological Conservation, 220, 1–11. https://doi.org/10.1016/j.biocon.2018.01.030

Watanabe, S. (2010). Asymptotic Equivalence of Bayes Cross Validation and Widely Applicable Information Criterion in Singular Learning Theory. The Journal of Machine Learning Research, 11, 3571–3594.

Watanabe, S. (2013). A Widely Applicable Bayesian Information Criterion. The Journal of Machine Learning Research, 14, 867–897.

Westfall, K. M., Therriault, T. W., & Abbott, C. L. (2019). A new approach to molecular biosurveillance of invasive species using DNA metabarcoding. Global Change Biology, n/a(n/a). https://doi.org/10.1111/gcb.14886

Yin, D., & He, F. (2014). A simple method for estimating species abundance from occurrence maps. Methods in Ecology and Evolution, 5(4), 336–343. https://doi.org/10.1111/2041-210X.12159

