## Supplementary FIgure 1 for "Targeted Next Generation Sequencing of environmental DNA improves detection and quantification of invasive European green crab (*Carcinus maenas*)"

**Supplementary Fig. 1** Predicted catch per unit effort for replicate data sets of the (a) tNGS and (b) qPCR assays. Grey surround is 95% confidence interval. Raw data shown in grey dots. Adjusted R^2^ values are (a) 0.864 and (b) 0.480.

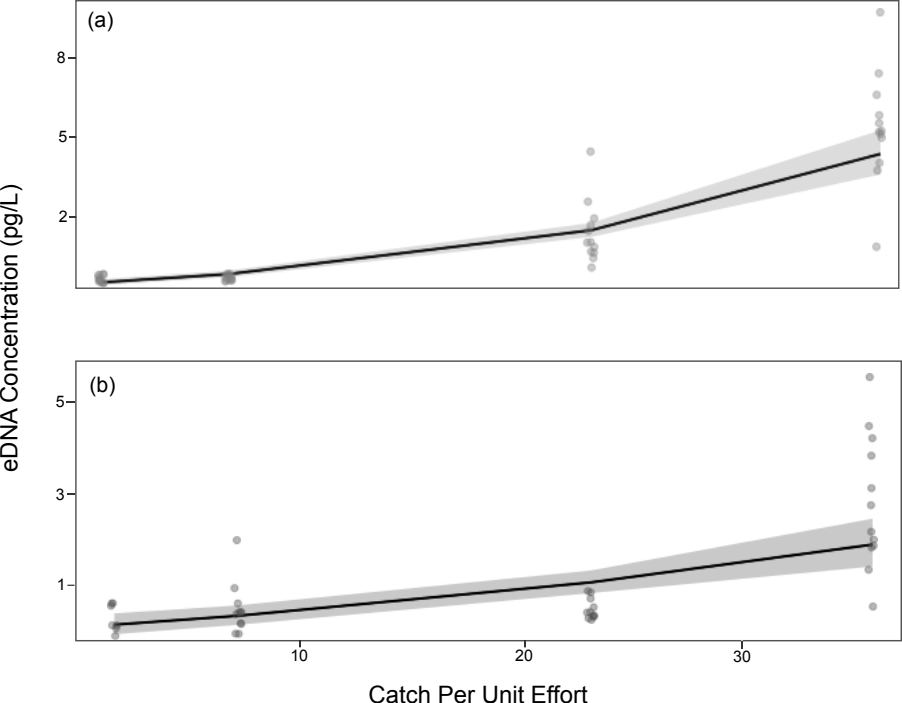
