## Supplementary Table 1 for "Targeted Next Generation Sequencing of environmental DNA improves detection and quantification of invasive European green crab (*Carcinus maenas*)"

**Supplementary Table S1** Geographical information, CPUE and relative abundance category for each site, refer to Fig. 1 for map. Minimum, maximum and SD among technical PCR replicates for qPCR C_t_ values for the *site* data set (above) and technical PCR replicates averaged among site replicates for the *replicate* data set (below). Minimum, maximum and SD of raw sequencing reads among site replicates for the *replicate* data set (below) and one value reported for the *site* data set (above). C_t_ = U is undetected.

|  |  |  |  |  |  | *C_t_* | | | *Number of green crab reads* | | |
| --- | --- | --- | --- | --- | --- | --- | --- | --- | --- | --- | --- |
| **Site** | **Latitude** | **Longitude** | **CPUE** | **Density** | **Day** | **MIN** | **MAX** | **SD** |  |  |  |
| Mayne Bay | 48.991 | -125.296 | 23.0 | high | 1 | 36.46 | 36.49 | 0.02 |  | 4.86E+05 |  |
|  |  |  |  |  | 2 | 36.28 | 36.29 | 0.01 |  | 1.06E+05 |  |
| Hillier Island | 49.029 | -125.320 | 73.5 | high | 1 | 37.34 | 37.34 | - |  | 5.06E+04 |  |
|  |  |  |  |  | 2 | 34.44 | 34.95 | 0.29 |  | 4.60E+05 |  |
| Pipestem Inlet | 49.030 | -125.251 | 35.4 | high | 1 | 34.27 | 34.61 | 0.18 |  | 6.21E+05 |  |
|  |  |  |  |  | 2 | 33.19 | 34.62 | 0.75 |  | 6.99E+05 |  |
| Ritherdon Bay | 48.951 | -124.981 | 7.4 | low | 1 | U | U | U |  | 1.10E+01 |  |
|  |  |  |  |  | 2 | 35.11 | 36.38 | 0.9 |  | 3.00E+01 |  |
| San Mateo Bay | 49.938 | -124.988 | 2.0 | low | 1 | 35.17 | 35.17 | - |  | 5.56E+04 |  |
|  |  |  |  |  | 2 | U | U | U |  | 1.85E+04 |  |
|  |  |  |  |  |  |  |  |  | **MIN** | **MAX** | **SD** |
| Mayne Bay |  |  |  |  | 1 | 34.20 | 38.55 | 1.23 | 2.74E+05 | 7.87E+05 | 1.98E+05 |
|  |  |  |  |  | 2 | 35.49 | 37.57 | 0.72 | 4.10E+04 | 4.11E+05 | 1.43E+05 |
| Pipestem Inlet |  |  |  |  | 1 | 33.49 | 37.83 | 1.16 | 2.55E+05 | 6.50E+05 | 1.54E+05 |
|  |  |  |  |  | 2 | 32.53 | 34.51 | 0.61 | 1.21E+05 | 7.51E+05 | 2.33E+05 |
| Ritherdon Bay |  |  |  |  | 1 | 35.48 | 38.88 | 1.08 | 5.00E+00 | 1.62E+04 | 6.60E+03 |
|  |  |  |  |  | 2 | 35.19 | 37.88 | 0.93 | 1.10E+01 | 5.74E+03 | 2.26E+03 |
| San Mateo Bay |  |  |  |  | 1 | 35.38 | 38.09 | 0.79 | 0.00E+00 | 2.30E+01 | 8.42E+00 |
|  |  |  |  |  | 2 | 35.92 | 37.75 | 1.05 | 2.00E+00 | 6.80E+01 | 2.83E+01 |
