## Supplementary Table 2 for "Targeted Next Generation Sequencing of environmental DNA improves detection and quantification of invasive European green crab (*Carcinus maenas*)"

**Supplementary Table S2** Sequences for fusion primers (5’ - [N5 + Sequencing Primer 1 + Inline Barcode + Base Shift N + Green Crab Primer] – 3’) & (5’ - [N7 + Sequencing Primer 2 + Inline Barcode + Base Shift N + Green Crab Primer] - 3’). Primer names containing ‘F’ are forward green crab primer, primer names containing ‘R’ are reverse green crab primer. Pairs are numbered 1-50.

| Name | N5 | Sequencing Primer 1 | Inline Barcode | Base shift N | Green Crab Primer | Pair |
| --- | --- | --- | --- | --- | --- | --- |
| N5_gcF1 | AATGATACGGCGACCACCGAGATCTACAC | TCTTTCCCTACACGACGCTCTTCCGATCT | AACCG |  | ATGAACAGTCTATCCTCCTTTAG | 1 |
| N5_gcF2 | AATGATACGGCGACCACCGAGATCTACAC | TCTTTCCCTACACGACGCTCTTCCGATCT | AACGC | C | ATGAACAGTCTATCCTCCTTTAG | 2 |
| N5_gcF3 | AATGATACGGCGACCACCGAGATCTACAC | TCTTTCCCTACACGACGCTCTTCCGATCT | TAATC | GA | ATGAACAGTCTATCCTCCTTTAG | 3 |
| N5_gcF4 | AATGATACGGCGACCACCGAGATCTACAC | TCTTTCCCTACACGACGCTCTTCCGATCT | TAAGA | TGC | ATGAACAGTCTATCCTCCTTTAG | 4 |
| N5_gcF5 | AATGATACGGCGACCACCGAGATCTACAC | TCTTTCCCTACACGACGCTCTTCCGATCT | AACTT | ACGC | ATGAACAGTCTATCCTCCTTTAG | 5 |
| N5_gcF6 | AATGATACGGCGACCACCGAGATCTACAC | TCTTTCCCTACACGACGCTCTTCCGATCT | AAGAC |  | ATGAACAGTCTATCCTCCTTTAG | 6 |
| N5_gcF7 | AATGATACGGCGACCACCGAGATCTACAC | TCTTTCCCTACACGACGCTCTTCCGATCT | TACAG | G | ATGAACAGTCTATCCTCCTTTAG | 7 |
| N5_gcF8 | AATGATACGGCGACCACCGAGATCTACAC | TCTTTCCCTACACGACGCTCTTCCGATCT | AAGCA | TC | ATGAACAGTCTATCCTCCTTTAG | 8 |
| N5_gcF9 | AATGATACGGCGACCACCGAGATCTACAC | TCTTTCCCTACACGACGCTCTTCCGATCT | ATGCC | AGC | ATGAACAGTCTATCCTCCTTTAG | 9 |
| N5_gcF10 | AATGATACGGCGACCACCGAGATCTACAC | TCTTTCCCTACACGACGCTCTTCCGATCT | AAGGT | TGCA | ATGAACAGTCTATCCTCCTTTAG | 10 |
| N5_gcF11 | AATGATACGGCGACCACCGAGATCTACAC | TCTTTCCCTACACGACGCTCTTCCGATCT | AAGTG |  | ATGAACAGTCTATCCTCCTTTAG | 11 |
| N5_gcF12 | AATGATACGGCGACCACCGAGATCTACAC | TCTTTCCCTACACGACGCTCTTCCGATCT | TAACT | T | ATGAACAGTCTATCCTCCTTTAG | 12 |
| N5_gcF13 | AATGATACGGCGACCACCGAGATCTACAC | TCTTTCCCTACACGACGCTCTTCCGATCT | TCGTT | AG | ATGAACAGTCTATCCTCCTTTAG | 13 |
| N5_gcF14 | AATGATACGGCGACCACCGAGATCTACAC | TCTTTCCCTACACGACGCTCTTCCGATCT | AATAG | CTG | ATGAACAGTCTATCCTCCTTTAG | 14 |
| N5_gcF15 | AATGATACGGCGACCACCGAGATCTACAC | TCTTTCCCTACACGACGCTCTTCCGATCT | TAGAT | GTAC | ATGAACAGTCTATCCTCCTTTAG | 15 |
| N5_gcF16 | AATGATACGGCGACCACCGAGATCTACAC | TCTTTCCCTACACGACGCTCTTCCGATCT | AATCC |  | ATGAACAGTCTATCCTCCTTTAG | 16 |
| N5_gcF17 | AATGATACGGCGACCACCGAGATCTACAC | TCTTTCCCTACACGACGCTCTTCCGATCT | TAGCC | A | ATGAACAGTCTATCCTCCTTTAG | 17 |
| N5_gcF18 | AATGATACGGCGACCACCGAGATCTACAC | TCTTTCCCTACACGACGCTCTTCCGATCT | AATGA | CT | ATGAACAGTCTATCCTCCTTTAG | 18 |
| N5_gcF19 | AATGATACGGCGACCACCGAGATCTACAC | TCTTTCCCTACACGACGCTCTTCCGATCT | TAGTA | GAG | ATGAACAGTCTATCCTCCTTTAG | 19 |
| N5_gcF20 | AATGATACGGCGACCACCGAGATCTACAC | TCTTTCCCTACACGACGCTCTTCCGATCT | ACAAC | CTTG | ATGAACAGTCTATCCTCCTTTAG | 20 |
| N5_gcF21 | AATGATACGGCGACCACCGAGATCTACAC | TCTTTCCCTACACGACGCTCTTCCGATCT | TATAA |  | ATGAACAGTCTATCCTCCTTTAG | 21 |
| N5_gcF22 | AATGATACGGCGACCACCGAGATCTACAC | TCTTTCCCTACACGACGCTCTTCCGATCT | ACACA | G | ATGAACAGTCTATCCTCCTTTAG | 22 |
| N5_gcF23 | AATGATACGGCGACCACCGAGATCTACAC | TCTTTCCCTACACGACGCTCTTCCGATCT | ACAGG | TT | ATGAACAGTCTATCCTCCTTTAG | 23 |
| N5_gcF24 | AATGATACGGCGACCACCGAGATCTACAC | TCTTTCCCTACACGACGCTCTTCCGATCT | TATGC | TGA | ATGAACAGTCTATCCTCCTTTAG | 24 |
| N5_gcF25 | AATGATACGGCGACCACCGAGATCTACAC | TCTTTCCCTACACGACGCTCTTCCGATCT | ACATT | AGTT | ATGAACAGTCTATCCTCCTTTAG | 25 |
| N5_gcR1 | AATGATACGGCGACCACCGAGATCTACAC | TCTTTCCCTACACGACGCTCTTCCGATCT | CCTTG |  | GAAAGAACGCATATTGATAATAGTTG | 26 |
| N5_gcR2 | AATGATACGGCGACCACCGAGATCTACAC | TCTTTCCCTACACGACGCTCTTCCGATCT | CGACT | A | GAAAGAACGCATATTGATAATAGTTG | 27 |
| N5_gcR3 | AATGATACGGCGACCACCGAGATCTACAC | TCTTTCCCTACACGACGCTCTTCCGATCT | CGATA | TC | GAAAGAACGCATATTGATAATAGTTG | 28 |
| N5_gcR4 | AATGATACGGCGACCACCGAGATCTACAC | TCTTTCCCTACACGACGCTCTTCCGATCT | CGCAA | CAC | GAAAGAACGCATATTGATAATAGTTG | 29 |
| N5_gcR5 | AATGATACGGCGACCACCGAGATCTACAC | TCTTTCCCTACACGACGCTCTTCCGATCT | CGCTT | CTAC | GAAAGAACGCATATTGATAATAGTTG | 30 |
| N5_gcR6 | AATGATACGGCGACCACCGAGATCTACAC | TCTTTCCCTACACGACGCTCTTCCGATCT | CGTAC |  | GAAAGAACGCATATTGATAATAGTTG | 31 |
| N5_gcR7 | AATGATACGGCGACCACCGAGATCTACAC | TCTTTCCCTACACGACGCTCTTCCGATCT | CGTCA | C | GAAAGAACGCATATTGATAATAGTTG | 32 |
| N5_gcR8 | AATGATACGGCGACCACCGAGATCTACAC | TCTTTCCCTACACGACGCTCTTCCGATCT | CTAAC | GT | GAAAGAACGCATATTGATAATAGTTG | 33 |
| N5_gcR9 | AATGATACGGCGACCACCGAGATCTACAC | TCTTTCCCTACACGACGCTCTTCCGATCT | GACTC | TGT | GAAAGAACGCATATTGATAATAGTTG | 34 |
| N5_gcR10 | AATGATACGGCGACCACCGAGATCTACAC | TCTTTCCCTACACGACGCTCTTCCGATCT | GAGAG | CACT | GAAAGAACGCATATTGATAATAGTTG | 35 |
| N5_gcR11 | AATGATACGGCGACCACCGAGATCTACAC | TCTTTCCCTACACGACGCTCTTCCGATCT | GAGGA |  | GAAAGAACGCATATTGATAATAGTTG | 36 |
| N5_gcR12 | AATGATACGGCGACCACCGAGATCTACAC | TCTTTCCCTACACGACGCTCTTCCGATCT | GAGTT | A | GAAAGAACGCATATTGATAATAGTTG | 37 |
| N5_gcR13 | AATGATACGGCGACCACCGAGATCTACAC | TCTTTCCCTACACGACGCTCTTCCGATCT | GATAC | AT | GAAAGAACGCATATTGATAATAGTTG | 38 |
| N5_gcR14 | AATGATACGGCGACCACCGAGATCTACAC | TCTTTCCCTACACGACGCTCTTCCGATCT | GATCA | GTA | GAAAGAACGCATATTGATAATAGTTG | 39 |
| N5_gcR15 | AATGATACGGCGACCACCGAGATCTACAC | TCTTTCCCTACACGACGCTCTTCCGATCT | GATGT | GTCA | GAAAGAACGCATATTGATAATAGTTG | 40 |
| N5_gcR16 | AATGATACGGCGACCACCGAGATCTACAC | TCTTTCCCTACACGACGCTCTTCCGATCT | GCACT |  | GAAAGAACGCATATTGATAATAGTTG | 41 |
| N5_gcR17 | AATGATACGGCGACCACCGAGATCTACAC | TCTTTCCCTACACGACGCTCTTCCGATCT | GCATG | T | GAAAGAACGCATATTGATAATAGTTG | 42 |
| N5_gcR18 | AATGATACGGCGACCACCGAGATCTACAC | TCTTTCCCTACACGACGCTCTTCCGATCT | GCCTA | CA | GAAAGAACGCATATTGATAATAGTTG | 43 |
| N5_gcR19 | AATGATACGGCGACCACCGAGATCTACAC | TCTTTCCCTACACGACGCTCTTCCGATCT | GCGAT | CTA | GAAAGAACGCATATTGATAATAGTTG | 44 |
| N5_gcR20 | AATGATACGGCGACCACCGAGATCTACAC | TCTTTCCCTACACGACGCTCTTCCGATCT | GCTAG | TGTA | GAAAGAACGCATATTGATAATAGTTG | 45 |
| N5_gcR21 | AATGATACGGCGACCACCGAGATCTACAC | TCTTTCCCTACACGACGCTCTTCCGATCT | GGAAT |  | GAAAGAACGCATATTGATAATAGTTG | 46 |
| N5_gcR22 | AATGATACGGCGACCACCGAGATCTACAC | TCTTTCCCTACACGACGCTCTTCCGATCT | GGATC | C | GAAAGAACGCATATTGATAATAGTTG | 47 |
| N5_gcR23 | AATGATACGGCGACCACCGAGATCTACAC | TCTTTCCCTACACGACGCTCTTCCGATCT | GGTGA | AC | GAAAGAACGCATATTGATAATAGTTG | 48 |
| N5_gcR24 | AATGATACGGCGACCACCGAGATCTACAC | TCTTTCCCTACACGACGCTCTTCCGATCT | GGTTG | TAA | GAAAGAACGCATATTGATAATAGTTG | 49 |
| N5_gcR25 | AATGATACGGCGACCACCGAGATCTACAC | TCTTTCCCTACACGACGCTCTTCCGATCT | GTAAG | ATCA | GAAAGAACGCATATTGATAATAGTTG | 50 |
|  | N7 | Sequencing Primer 2 | Inline Barcode | Base shift N | Green Crab Primer | Pair |
| N7_gcR1 | CAAGCAGAAGACGGCATACGAGAT | CGGTCTCGGCATTCCTGCTGAACCGCTCTTCCGATCT | CCTTG |  | GAAAGAACGCATATTGATAATAGTTG | 5 |
| N7_gcR2 | CAAGCAGAAGACGGCATACGAGAT | CGGTCTCGGCATTCCTGCTGAACCGCTCTTCCGATCT | CGACT | A | GAAAGAACGCATATTGATAATAGTTG | 4 |
| N7_gcR3 | CAAGCAGAAGACGGCATACGAGAT | CGGTCTCGGCATTCCTGCTGAACCGCTCTTCCGATCT | CGATA | TC | GAAAGAACGCATATTGATAATAGTTG | 3 |
| N7_gcR4 | CAAGCAGAAGACGGCATACGAGAT | CGGTCTCGGCATTCCTGCTGAACCGCTCTTCCGATCT | CGCAA | CAC | GAAAGAACGCATATTGATAATAGTTG | 2 |
| N7_gcR5 | CAAGCAGAAGACGGCATACGAGAT | CGGTCTCGGCATTCCTGCTGAACCGCTCTTCCGATCT | CGCTT | CTAC | GAAAGAACGCATATTGATAATAGTTG | 1 |
| N7_gcR6 | CAAGCAGAAGACGGCATACGAGAT | CGGTCTCGGCATTCCTGCTGAACCGCTCTTCCGATCT | CGTAC |  | GAAAGAACGCATATTGATAATAGTTG | 10 |
| N7_gcR7 | CAAGCAGAAGACGGCATACGAGAT | CGGTCTCGGCATTCCTGCTGAACCGCTCTTCCGATCT | CGTCA | C | GAAAGAACGCATATTGATAATAGTTG | 9 |
| N7_gcR8 | CAAGCAGAAGACGGCATACGAGAT | CGGTCTCGGCATTCCTGCTGAACCGCTCTTCCGATCT | CTAAC | GT | GAAAGAACGCATATTGATAATAGTTG | 8 |
| N7_gcR9 | CAAGCAGAAGACGGCATACGAGAT | CGGTCTCGGCATTCCTGCTGAACCGCTCTTCCGATCT | GACTC | TGT | GAAAGAACGCATATTGATAATAGTTG | 7 |
| N7_gcR10 | CAAGCAGAAGACGGCATACGAGAT | CGGTCTCGGCATTCCTGCTGAACCGCTCTTCCGATCT | GAGAG | CACT | GAAAGAACGCATATTGATAATAGTTG | 6 |
| N7_gcR11 | CAAGCAGAAGACGGCATACGAGAT | CGGTCTCGGCATTCCTGCTGAACCGCTCTTCCGATCT | GAGGA |  | GAAAGAACGCATATTGATAATAGTTG | 15 |
| N7_gcR12 | CAAGCAGAAGACGGCATACGAGAT | CGGTCTCGGCATTCCTGCTGAACCGCTCTTCCGATCT | GAGTT | A | GAAAGAACGCATATTGATAATAGTTG | 14 |
| N7_gcR13 | CAAGCAGAAGACGGCATACGAGAT | CGGTCTCGGCATTCCTGCTGAACCGCTCTTCCGATCT | GATAC | AT | GAAAGAACGCATATTGATAATAGTTG | 13 |
| N7_gcR14 | CAAGCAGAAGACGGCATACGAGAT | CGGTCTCGGCATTCCTGCTGAACCGCTCTTCCGATCT | GATCA | GTA | GAAAGAACGCATATTGATAATAGTTG | 12 |
| N7_gcR15 | CAAGCAGAAGACGGCATACGAGAT | CGGTCTCGGCATTCCTGCTGAACCGCTCTTCCGATCT | GATGT | GTCA | GAAAGAACGCATATTGATAATAGTTG | 11 |
| N7_gcR16 | CAAGCAGAAGACGGCATACGAGAT | CGGTCTCGGCATTCCTGCTGAACCGCTCTTCCGATCT | GCACT |  | GAAAGAACGCATATTGATAATAGTTG | 20 |
| N7_gcR17 | CAAGCAGAAGACGGCATACGAGAT | CGGTCTCGGCATTCCTGCTGAACCGCTCTTCCGATCT | GCATG | T | GAAAGAACGCATATTGATAATAGTTG | 19 |
| N7_gcR18 | CAAGCAGAAGACGGCATACGAGAT | CGGTCTCGGCATTCCTGCTGAACCGCTCTTCCGATCT | GCCTA | CA | GAAAGAACGCATATTGATAATAGTTG | 18 |
| N7_gcR19 | CAAGCAGAAGACGGCATACGAGAT | CGGTCTCGGCATTCCTGCTGAACCGCTCTTCCGATCT | GCGAT | CTA | GAAAGAACGCATATTGATAATAGTTG | 17 |
| N7_gcR20 | CAAGCAGAAGACGGCATACGAGAT | CGGTCTCGGCATTCCTGCTGAACCGCTCTTCCGATCT | GCTAG | TGTA | GAAAGAACGCATATTGATAATAGTTG | 16 |
| N7_gcR21 | CAAGCAGAAGACGGCATACGAGAT | CGGTCTCGGCATTCCTGCTGAACCGCTCTTCCGATCT | GGAAT |  | GAAAGAACGCATATTGATAATAGTTG | 25 |
| N7_gcR22 | CAAGCAGAAGACGGCATACGAGAT | CGGTCTCGGCATTCCTGCTGAACCGCTCTTCCGATCT | GGATC | C | GAAAGAACGCATATTGATAATAGTTG | 24 |
| N7_gcR23 | CAAGCAGAAGACGGCATACGAGAT | CGGTCTCGGCATTCCTGCTGAACCGCTCTTCCGATCT | GGTGA | AC | GAAAGAACGCATATTGATAATAGTTG | 23 |
| N7_gcR24 | CAAGCAGAAGACGGCATACGAGAT | CGGTCTCGGCATTCCTGCTGAACCGCTCTTCCGATCT | GGTTG | TAA | GAAAGAACGCATATTGATAATAGTTG | 22 |
| N7_gcR25 | CAAGCAGAAGACGGCATACGAGAT | CGGTCTCGGCATTCCTGCTGAACCGCTCTTCCGATCT | GTAAG | ATCA | GAAAGAACGCATATTGATAATAGTTG | 21 |
| N7_gcF1 | CAAGCAGAAGACGGCATACGAGAT | CGGTCTCGGCATTCCTGCTGAACCGCTCTTCCGATCT | AACCG |  | ATGAACAGTCTATCCTCCTTTAG | 30 |
| N7_gcF2 | CAAGCAGAAGACGGCATACGAGAT | CGGTCTCGGCATTCCTGCTGAACCGCTCTTCCGATCT | AACGC | C | ATGAACAGTCTATCCTCCTTTAG | 29 |
| N7_gcF3 | CAAGCAGAAGACGGCATACGAGAT | CGGTCTCGGCATTCCTGCTGAACCGCTCTTCCGATCT | TAATC | GA | ATGAACAGTCTATCCTCCTTTAG | 28 |
| N7_gcF4 | CAAGCAGAAGACGGCATACGAGAT | CGGTCTCGGCATTCCTGCTGAACCGCTCTTCCGATCT | TAAGA | TGC | ATGAACAGTCTATCCTCCTTTAG | 27 |
| N7_gcF5 | CAAGCAGAAGACGGCATACGAGAT | CGGTCTCGGCATTCCTGCTGAACCGCTCTTCCGATCT | AACTT | ACGC | ATGAACAGTCTATCCTCCTTTAG | 26 |
| N7_gcF6 | CAAGCAGAAGACGGCATACGAGAT | CGGTCTCGGCATTCCTGCTGAACCGCTCTTCCGATCT | AAGAC |  | ATGAACAGTCTATCCTCCTTTAG | 35 |
| N7_gcF7 | CAAGCAGAAGACGGCATACGAGAT | CGGTCTCGGCATTCCTGCTGAACCGCTCTTCCGATCT | TACAG | G | ATGAACAGTCTATCCTCCTTTAG | 34 |
| N7_gcF8 | CAAGCAGAAGACGGCATACGAGAT | CGGTCTCGGCATTCCTGCTGAACCGCTCTTCCGATCT | AAGCA | TC | ATGAACAGTCTATCCTCCTTTAG | 33 |
| N7_gcF9 | CAAGCAGAAGACGGCATACGAGAT | CGGTCTCGGCATTCCTGCTGAACCGCTCTTCCGATCT | ATGCC | AGC | ATGAACAGTCTATCCTCCTTTAG | 32 |
| N7_gcF10 | CAAGCAGAAGACGGCATACGAGAT | CGGTCTCGGCATTCCTGCTGAACCGCTCTTCCGATCT | AAGGT | TGCA | ATGAACAGTCTATCCTCCTTTAG | 31 |
| N7_gcF11 | CAAGCAGAAGACGGCATACGAGAT | CGGTCTCGGCATTCCTGCTGAACCGCTCTTCCGATCT | AAGTG |  | ATGAACAGTCTATCCTCCTTTAG | 40 |
| N7_gcF12 | CAAGCAGAAGACGGCATACGAGAT | CGGTCTCGGCATTCCTGCTGAACCGCTCTTCCGATCT | TAACT | T | ATGAACAGTCTATCCTCCTTTAG | 39 |
| N7_gcF13 | CAAGCAGAAGACGGCATACGAGAT | CGGTCTCGGCATTCCTGCTGAACCGCTCTTCCGATCT | TCGTT | AG | ATGAACAGTCTATCCTCCTTTAG | 38 |
| N7_gcF14 | CAAGCAGAAGACGGCATACGAGAT | CGGTCTCGGCATTCCTGCTGAACCGCTCTTCCGATCT | AATAG | CTG | ATGAACAGTCTATCCTCCTTTAG | 37 |
| N7_gcF15 | CAAGCAGAAGACGGCATACGAGAT | CGGTCTCGGCATTCCTGCTGAACCGCTCTTCCGATCT | TAGAT | GTAC | ATGAACAGTCTATCCTCCTTTAG | 36 |
| N7_gcF16 | CAAGCAGAAGACGGCATACGAGAT | CGGTCTCGGCATTCCTGCTGAACCGCTCTTCCGATCT | AATCC |  | ATGAACAGTCTATCCTCCTTTAG | 45 |
| N7_gcF17 | CAAGCAGAAGACGGCATACGAGAT | CGGTCTCGGCATTCCTGCTGAACCGCTCTTCCGATCT | TAGCC | A | ATGAACAGTCTATCCTCCTTTAG | 44 |
| N7_gcF18 | CAAGCAGAAGACGGCATACGAGAT | CGGTCTCGGCATTCCTGCTGAACCGCTCTTCCGATCT | AATGA | CT | ATGAACAGTCTATCCTCCTTTAG | 43 |
| N7_gcF19 | CAAGCAGAAGACGGCATACGAGAT | CGGTCTCGGCATTCCTGCTGAACCGCTCTTCCGATCT | TAGTA | GAG | ATGAACAGTCTATCCTCCTTTAG | 42 |
| N7_gcF20 | CAAGCAGAAGACGGCATACGAGAT | CGGTCTCGGCATTCCTGCTGAACCGCTCTTCCGATCT | ACAAC | CTTG | ATGAACAGTCTATCCTCCTTTAG | 41 |
| N7_gcF21 | CAAGCAGAAGACGGCATACGAGAT | CGGTCTCGGCATTCCTGCTGAACCGCTCTTCCGATCT | TATAA |  | ATGAACAGTCTATCCTCCTTTAG | 50 |
| N7_gcF22 | CAAGCAGAAGACGGCATACGAGAT | CGGTCTCGGCATTCCTGCTGAACCGCTCTTCCGATCT | ACACA | G | ATGAACAGTCTATCCTCCTTTAG | 49 |
| N7_gcF23 | CAAGCAGAAGACGGCATACGAGAT | CGGTCTCGGCATTCCTGCTGAACCGCTCTTCCGATCT | ACAGG | TT | ATGAACAGTCTATCCTCCTTTAG | 48 |
| N7_gcF24 | CAAGCAGAAGACGGCATACGAGAT | CGGTCTCGGCATTCCTGCTGAACCGCTCTTCCGATCT | TATGC | TGA | ATGAACAGTCTATCCTCCTTTAG | 47 |
| N7_gcF25 | CAAGCAGAAGACGGCATACGAGAT | CGGTCTCGGCATTCCTGCTGAACCGCTCTTCCGATCT | ACATT | AGTT | ATGAACAGTCTATCCTCCTTTAG | 46 |
