## Supplementary Table 3 for "Targeted Next Generation Sequencing of environmental DNA improves detection and quantification of invasive European green crab (*Carcinus maenas*)"

**Supplementary Table S3** Posterior means (parameter estimates) and model selection criteria (PPLC = Posterior Predictive Loss Criterion and WAIC = Widely Accepted Information Criterion) for Bayesian occupancy models fitted to green crab eDNA detection results. Models with and without catch per unit effort (CPUE) as a covariate were run for 33,000 (tNGS) and 11,000 (qPCR) MCMC iterations and parameters estimated after discarding 10% burnin runs. Selected model is in bold for each assay.

| Assay | Model | Occupancy at Site (*ψ*) | Occupancy in Sample (*α*) | Detection in Replicate (*ρ*) | PPLC | WAIC |
| --- | --- | --- | --- | --- | --- | --- |
| tNGS | *ψ*(•), *θ*(•), *ρ*(•) | 0.87 | (1.591)′ | 0.94 | 5.25 | 14.98 |
| tNGS | ***ψ*(•), *θ*(CPUE), *ρ*(•)** | 0.87 | (1,657,0.799)′ | 0.97 | 2.78 | 9.75 |
| qPCR | *ψ*(•), *θ*(•), *ρ*(•) | 0.87 | (1.096)′ | 0.74 | 50.74 | 56.16 |
| qPCR | ***ψ*(•), *θ*(CPUE), *ρ*(•)** | 0.87 | (1.368,0.857)′ | 0.74 | 49.47 | 52.85 |
